# Changes in Secondary Structure and Properties of Bovine Serum Albumin as a Result of Interactions with Gold Surface

**DOI:** 10.1101/2023.11.18.567678

**Authors:** Paulina Komorek, Kamil Rakowski, Magdalena Szota, Małgorzata Lekka, Barbara Jachimska

## Abstract

Proteins can alter their shape when interacting with a surface. This study explores how bovine serum albumin (BSA) modifies structurally when it adheres to a gold surface, depending on the protein concentration and pH. We verified that the gold surface induces significant structural modifications to the BSA molecule using circular dichroism, infrared spectroscopy, and atomic force microscopy. Specifically, adsorbed molecules displayed increased levels of disordered structures and β-turns, with fewer α-helices than the native structure. MP-SPR spectroscopy demonstrated that the protein molecules preferred a planar orientation during adsorption. Molecular dynamics simulations revealed that the interaction between cysteines exposed to the outside of the molecule and the gold surface was vital, especially at pH = 3.5. The macroscopic properties of the protein film observed by AFM and contact angles confirm the flexible nature of the protein itself. Notably, structural transformation is joined with the degree of hydration of protein layers.

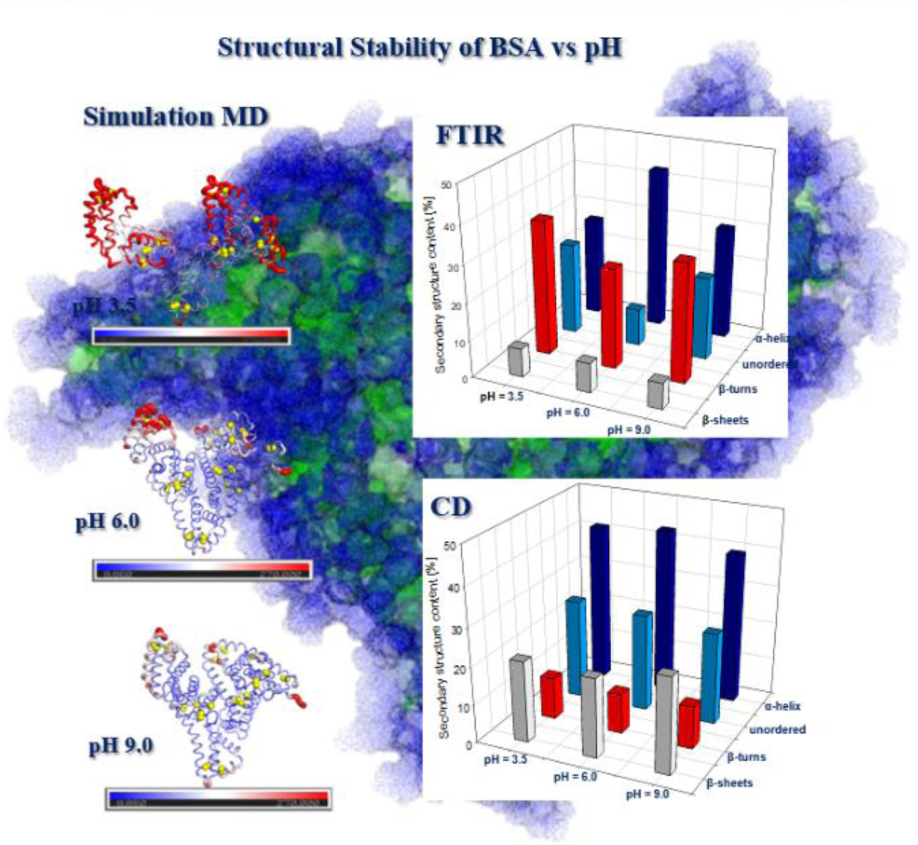

## Introduction

Application of proteins in biomedical areas requires controlling the conformation of the molecules, which is directly related to the activity of proteins and, thus, their biological functions [1]. Among the many parameters that affect the conformation of proteins, the impact of interactions with the surface is much less understood than factors such as pH, solvent type, or temperature [2–12]. Bovine serum albumin (BSA) is widely studied as a model globular protein [13][14]. Additionally, the popularity of BSA results from its ability to bind ions and various types of molecules, such as drugs, with blood composition regulation being its biological function [15][16]. The BSA molecule comprises 583 amino acids, with a molar mass of 66,430 Da. BSA is a single chain cross-linked with 17 disulfide bridges and one thiol bridge [17][18]. The molecule is heart-shaped and comprises 3 homologous but structurally distinct domains (I, II, and III), which are stabilized by an internal network of disulfide bonds and each bearing many ionizable groups with opposite signs [18][19]. Many studies have been conducted using various spectroscopic techniques to explore changes in the BSA structure dependent on environmental conditions. Performed studies using FTIR, CD, and NMR showed that BSA could adopt three main conformations (N-form, F-form, E-form) depending on the pH of the solvent. The loss of BSA stability and the expansion of the structure progresses with decreasing pH, where under acidic conditions, the protein structure becomes increasingly elongated and is characterized by marked decreases of alpha-helices [20 - 23]. Moreover, it was confirmed that conformational changes of BSA could be reversible in a wide pH range (pH = 2.5–10.2) [24]. Another factor having a strong influence on the BSA conformation is temperature. It was presented that conformational changes of the BSA molecule are reversible in the temperature range of 42°C to 50°C. However, increasing the temperature above 60°C leads to irreversible aggregation [25][26]. Furthermore, a well-known phenomenon is the misfolding of BSA resulting from its interactions with different types of surfactants, drug molecules, or denaturants (urea and glutamine of hydrochloric acid, metal ions) [24]. Many studies have been conducted on the interaction of BSA with the surface, but they focused primarily on the kinetics of adsorption, adsorption efficiency, or viscoelastic properties of the obtained protein films. Currently, there are limited available studies on structural changes of BSA as a result of interactions with the surface [27–31]. As can be seen, the complexity of BSA conformational transitions is strongly dependent on the physical conditions. Simulation methods with atomic accuracy may allow the approximation of complex conformational transitions of BSA to see the complexity of carbon chain transformations at the domain and subdomain level. Importantly, the mechanism of albumin’s interaction with the surface can be driven by the high affinity of disulfides and thiol groups for gold atoms, as demonstrated by experimental methods based on Raman spectroscopy [32][33]. Known factors affecting the conformational changes of albumin, such as pH, raise the question of how the accessibility of the disulfide chains to potential reductants such as gold surfaces will change. Therefore, this work attempted to determine the conformational effects of BSA and properties on obtained protein films due to the interaction with a solid surface. Depending on pH, changes in the secondary structure of BSA in bulk solution were determined using Circular Dichroism (CD). The BSA residues which are responsible for the greatest conformational changes were established by Molecular Dynamics simulations (MD). With protonation corresponding to acidic and alkaline conditions, the structural flexibility of BSA and the surface availability of cysteines were compared. In addition, based on the last frames of the simulation, hydrophobicity maps of the BSA surface were generated. The effect of interactions between the protein and gold surface was investigated by comparing the secondary structure obtained by CD with the structure adsorbed BSA molecules on the gold surface obtained by Fourier Transform Infrared Spectroscopy (FTIR). Multi-Parametric Surface Plasmon Resonance (MP-SPR) and Quartz Crystal Microbalance with Dissipation Monitoring (QCM-D) were used to estimate the effectiveness of BSA adsorption. The MP-SPR results, combined with the random sequential adsorption (RSA) model, were also used to determine the orientation of molecules in adsorbed films. Comparing the adsorbed mass obtained by the MP-SPR and QCM-D enabled films’ hydration calculations. The conclusions presented in this work may contribute to improvements in controlling the BSA molecules’ conformation and protein films’ structure used to form biomaterials for biomedical and nanotechnology applications.

## Materials and Methods

### Materials

Experiments were conducted using Bovine Serum Albumin (BSA) purchased at *Merck & Co*., NJ, USA. The purity of BSA was equal to 99%, and it was used without further purification. The BSA solutions with different protein concentrations were prepared by dissolving an appropriate amount of protein powder in sodium chloride (NaCl) solutions with two different ionic strengths (I = 0.01 M or I = 0.15 M). The pH of solutions was adjusted by adding small amounts of hydrochloric acid (HCl) or sodium hydroxide (NaOH) solutions. The pH of BSA and NaCl solutions was monitored using a precise pH meter (*WTW*, London, UK) equipped with an electrode (*Hamilton*, NA, USA).

### Methods

Atomic Force Microscopy (AFM) was used to characterize the topography of the protein films as a function of pH (pH = 4.0; 6.0 or 8.0). The experiments were carried out using the AFM instrument model XE-120 manufactured by *Park System* (CA, USA) working in the contact mode. Measurements were performed with MLCT-D AFM probes (*Bruker*, MA, USA) made of silicon nitride (Si_3_N_4_) covered with a layer of gold on the backside. The nominal parameters of the probes, such as the tip radius, the cantilever’s length, and width, were equal to 20 nm, 225 μm, and 20 μm, respectively. The AFM probe resonant frequency was 15 kHz, and a nominal spring constant equaled 0.03 N/m.

Multi-Parametric Surface Plasmon Resonance (MP-SPR) *NaviTM* 200 (*BioNavis*, Finland) apparatus with a goniometer coupled to a prism (Krechmer mode) was used to determine the efficiency of BSA adsorption on the gold surface. The experiments depended on BSA concentration (c = 5–200 ppm) or pH (pH = 4.0–9.0) for ionic strength I = 0.01 M or I = 0.15 M. The system has two separate channels, each emitting 670 and 785 nm wavelengths, and it operates in a wide range of scanning angles (40–78◦). A peristaltic pump is also included in the system. The measurements were taken as follows: a baseline for a NaCl solution with a defined ionic strength (I = 0.01 M or I = 0.15 M) and selected pH, then 90 min of BSA adsorption with a specified concentration and pH, and 90 min of rinsing with NaCl solution at the specified ionic strength and pH.

Quartz Crystal Microbalance with Dissipation Monitoring (QCM-D) E1 *Q-Sense* (*Biolin Scientific*, Finland) allows recording two parameters during experiments: frequency changes in the sensor’s vibrations (Δf) and alterations in dissipation of the sensor’s energy (ΔD). Based on the obtained ΔD values, the viscoelastic properties of the adsorbed BSA films on the gold surface were determined. Recorded Δf allowed analysis of BSA adsorption’s effectiveness and reversibility/irreversibility. Given that the ΔD values showed that formed films are rigid, the Sauerbrey model converted Δf into a mass of adsorbed BSA. No considerable differences were noted for successive overtones in the system under examination, so the results were presented only for one overtone (n = 7).

Circular Dichroism (CD) was used to analyze changes in the secondary structure of BSA in solution dependent on environmental conditions (pH = 2.0–10.5, I = 0.01 M). Experiments were conducted using the Jasco-1500 spectrometer (*Jasco*, MD, USA) with a 10 nm quartz cuvette. In all measurements, the concentration of BSA was 50 ppm. The spectra were recorded in the wavelength range 185–300 nm with a resolution of 1 nm, while the scanning speed was equal to 50 nm/min. Quantitative analysis of BSA secondary structure was carried out using *Jasco* software (MD, USA).

Molecular Dynamics (MD) simulations were performed with the GROMACS software (version 2022) [34] and CHARMM-GUI meta server (Lehigh University, version 3.8, PA, USA) [35]. Three BSA structures were built with different protonation states and the appropriate number of TIP3P water. Cl^−^ and Na^+^ ions were added to the simulation boxes with the CHARMM36m force field and temperature of 298.15K. The PDB2PQR Server (Massachusetts Institute of Technology, version 3.5.2, MA, USA) [36] allowed the establishment of the three BSA protonation states corresponding to pH = 3.5, pH = 6.0, or pH = 9.0. For simulations conducted at pH = 3.5, 69 ions were added, therein 61 counterions (Cl^−^); at pH = 6.0, 19 ions and 15 counterions (Na^+^), and at pH = 9.0–26 ions, therein 22 counterions (Na^+^) were included. In the simulations, the cubic cell was used: for pH = 3.5 with a side length of 180 Å, for pH 6.0 with a length of 120 Å, and for pH = 9.0, 130 Å. In pH = 6.0 and pH = 9.0, the initial protein edge distance was equal to 12 Å, and at pH = 3.5 system had the largest cell to ensure the transition of the BSA N-form and F-form. Energy minimization was performed with the fastest-decreasing algorithm with a time step of 5000 ps. The equilibration was conducted with the Nose-Hoover thermostat with the coupling constant τ = 1.0 ps and the Parrinello-Rahman barostat with a coupling constant τ = 5.0 ps (isotropic). The simulations were run for 200 ns. The Pymol (*Schrödinger, Inc*., version 2.5.4, USA, NY, USA) [37], BIOVIA Discovery Studio (*Dassault Systemes* BIOVIA, version 2021, CSA, USA) [38], and VMD (the University of Illinois Urbana-Champaign, version 1.9.4, IL, USA) [39] programs allowed for BSA states’ visualization. Mean square fluctuations in the BSA chain (RMSF) were studied to understand the influence of protonation on the local dynamics of essential parts of the protein chain. At the final section of the trajectory, where the RMSF was measured and in which the proteins reached the plateau of the RMSD function, ten models were generated, distant by the same time interval. From the structures obtained using the above method, the depth of cysteines from the surface accessible to the solvent was calculated, and the results were averaged and presented as a bar graph. The calculations were carried out analogously for each protonation state, and the depth of selected amino acids was determined by *EDTSurf* software (the University of Michigan, version 0.2009-10, MI, USA) [40]. Deformations of the BSA helices resulting from the N-F conformational transition were calculated by *Bendix* (VMD) algorithm. Values of dipole moments were determined from trajectories as a function of time using the Dipole Watcher (VMD) algorithm, and dipole moment vectors were visualized using the last frames.

Fourier Transform Infrared Spectroscopy (FTIR) measurements were carried out using a Nicolet iS10 spectrometer (*Thermo Fisher Scientific*, MA, USA) with the grazing angle-attenuated total reflection (VariGATR) accessory. The measurements were performed to investigate changes in BSA secondary structure due to protein interactions with the gold surface. A layer of gold with a thickness of 100 nm was deposited on the glass plate by vapor deposition. Measurements were recorded in the wavenumber range from 700–4000 cm^−1^. Overall, 128 scans were averaged for each spectrum with a spectral resolution of 4 1/cm. Before each measurement, the background spectrum was recorded and automatically subtracted from the spectrum of the protein sample. The measurements were carried out in varied pH conditions (pH = 2.0–11.0) and BSA concentration (c = 5–500 ppm). The adsorption time was 90 min. Afterward, the gold surface was rinsed successively for 90 min with 0.01 M NaCl solution with controlled pH. Finally, the surface was rinsed with water at the appropriate pH for 90 min. Before FTIR measurements, the gold surface with the adsorbed protein was dried with air. Omnic (*Thermo Fisher Scientific*, MA, USA) and Origin (*OriginLab*, MA, USA) software packages were used for data analysis.

Contact Angle (CA) measurements were performed using an axisymmetric drop shape analysis (ADSA) system with a system accuracy of 1°. For the CA measurements, BSA was adsorbed on the gold surface from a bulk solution (I = 0.01 M) with protein concentration c = 50 ppm in a wide pH range of 2.0−12.0. The BSA adsorption was carried out for 90 min, rinsing was kept for 90 min with I = 0.01 M NaCl solution with controlled pH and then the film was rinsed with water at the appropriate pH for 90 min. The image of a drop of sitting water was obtained with a CCD camera, and the drop’s shape on the sensor surface was fitted by the Young-Laplace equation.

## Results and Discussion

Conformational Stability of BSA in Bulk Solution Dependent on pH One of the main factors influencing the structure of BSA is the charge distribution on the molecule’s surface [27], which can be controlled by altering the pH. It was previously reported in the literature that there are three main conformational forms of the BSA molecule: elongated (E), fast (F), and native (N) [18]. The form in which the BSA molecule occurs depends on external factors. It has been observed that in a solution with a pH lower than 3.5, BSA is present in the E-form. At pH = 4.0, the molecules are in the N- and F-forms in equal proportions, while above pH = 4.5, the molecules are in their native state (N-form) [41]. To determine the composition of the BSA secondary structure dependent on pH, circular dichroism (CD) measurements were performed. According to the literature, BSA is a predominantly α-helical structure with several isomeric forms at different pH, corresponding to different α-helix content [27][42]. CD spectra of BSA as a function of pH are presented in Figure 1a. They are characterized by one positive band at 193 nm and two negative bands at 208 and 222 nm, characteristic of proteins with an α-helical structure [43][44]. Two negative peaks at 208 and 222 nm contribute to the π → π * and n → π* transfer for the α-helix peptide bond [45–47]. The presented results demonstrate slight changes in secondary structure in the pH range from 4.0 to 10.5. However, below pH = 4.0 a decrease in the intensity of the three main peaks is observed, related to the loss of α-helix content under acidic conditions. The CD spectra were used to probe BSA’s secondary structure, conformation, and stability in the solution. Obtained values of secondary structure components content dependent on pH are presented in Figure 1b. The measurement error was estimated by the measurement uncertainty, which was 5%. According to the results, BSA has a stable structure in a wide pH range (pH = 4.0-10.5) and its largest part consists of α-helices. Disordered structures achieve the second largest contribution in the total structure of BSA molecule, while the total content of β-sheets and β-turns is lower than 40%. The results are consistent with those obtained in the literature for BSA at pH = 7.0 [48][49]. Decreasing the pH below pH = 4.0 is associated with a change in the content of all the listed components of the BSA secondary structure. The contents of α-helices and β-sheets change the most, reaching 25% and 39%, respectively. The same tendency to decrease the content of α-helices and increase the content of β-sheets during BSA unfolding was also noticed in other research [50–52]. This process is related to rearranging the BSA secondary structure components adopting an E-form below pH = 3.5 [19].

**Figure 1.**
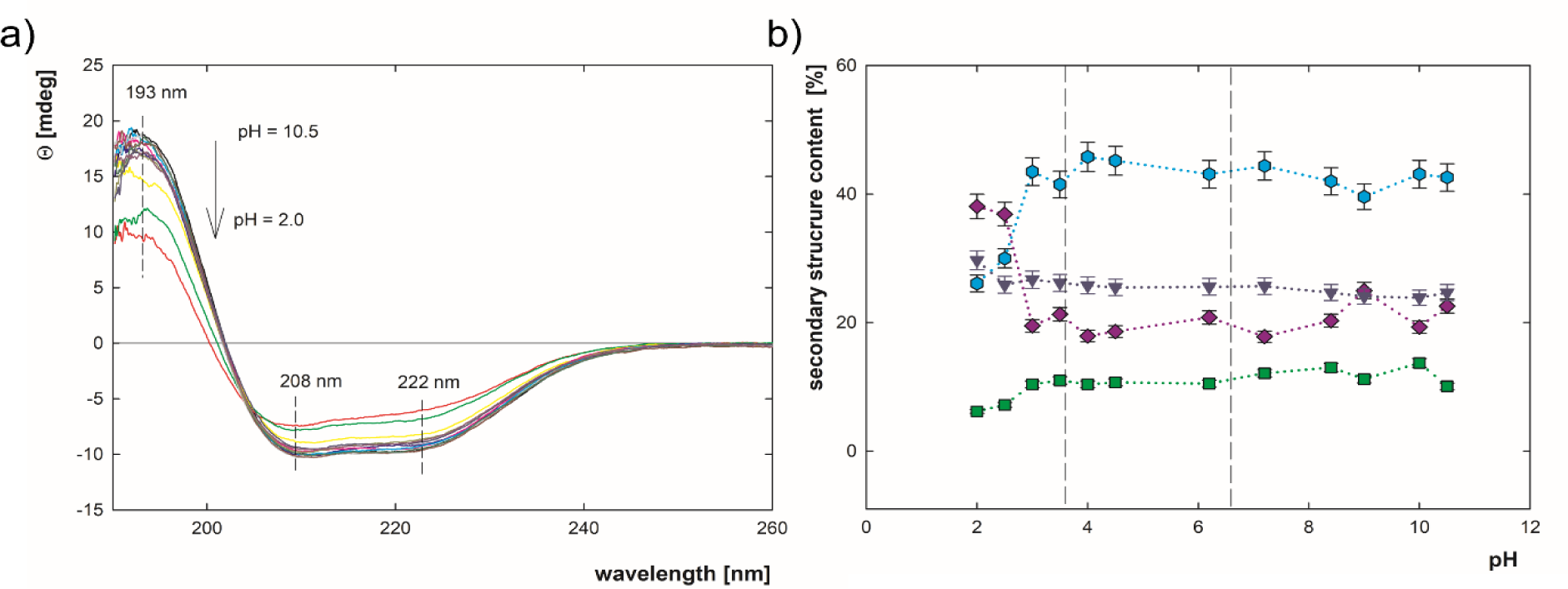
a) CD spectra for BSA, b) secondary structure components contribution dependent on pH (c = 50ppm, I = 0.01 M NaCl, pH = 2.0−10.5)

Measurements of the BSA secondary structure components in bulk solution showed that the BSA conformation is stable until pH = 4.0. However, in pH below 4.0, the BSA misfolding with simultaneous opening of its structure occurs [27].

### BSA Topography on the Gold Surface

Changes in the secondary structure of a protein may cause different surface protein topography. The topography of BSA on the surface may result not only from the differences in the BSA secondary structure taking place in the bulk solution depending on the pH but also due to the BSA-surface interactions, which may affect the structure of the adsorbed molecules. AFM was used to monitor the pH-dependent topography of BSA adsorbed on the gold surface. In the contact mode, the height of the soft structures is smaller due to the high pressure applied by the probing tip. Moreover, if the diameter of surface structures (such as proteins) is smaller than the radius of the probing tip, topographic images of such structures broaden. The resulting molecule’s shape and dimensions are the convolution of the AFM probe shape and the measured molecule [53]. For paraboloidal AFM probes, the true dimensions of the studied molecules a_k_ can be calculated from the following equation [54]:

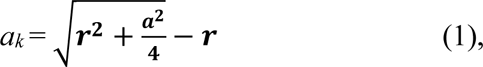

where *a* – the direct dimension of the structure measured by the AFM probe, *r* – tip radius of the paraboloidal AFM probe, here equal to 20 nm (the nominal value). The topographic images with the obtained true dimensions of the BSA molecules are presented in Table 1.

**Table 1.**
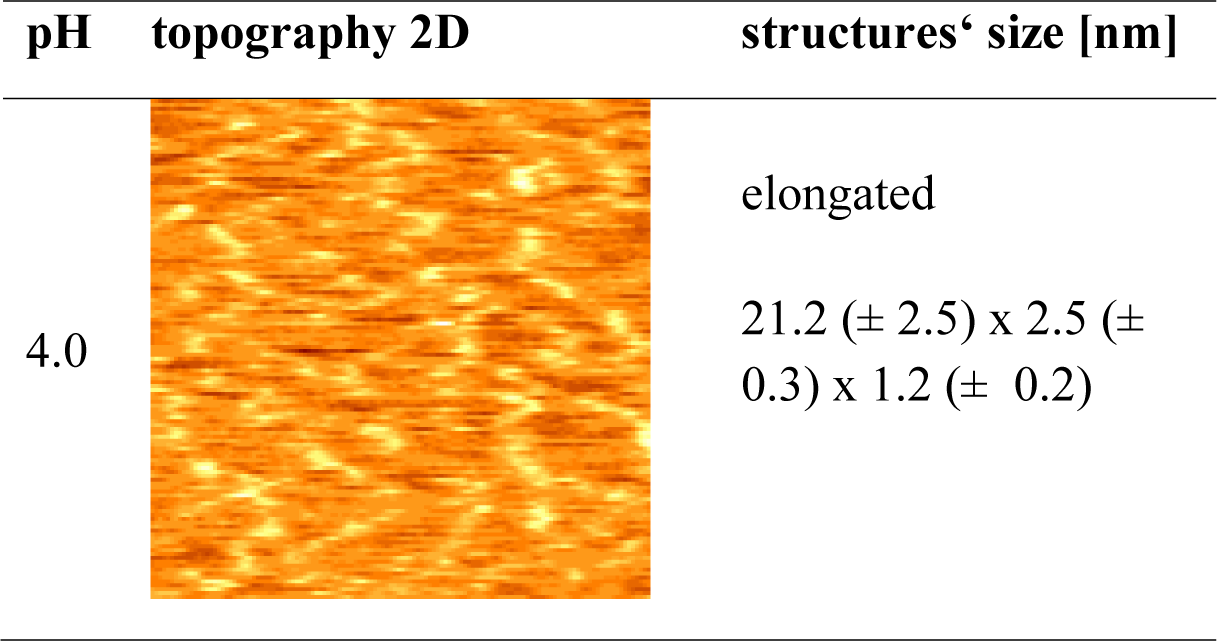
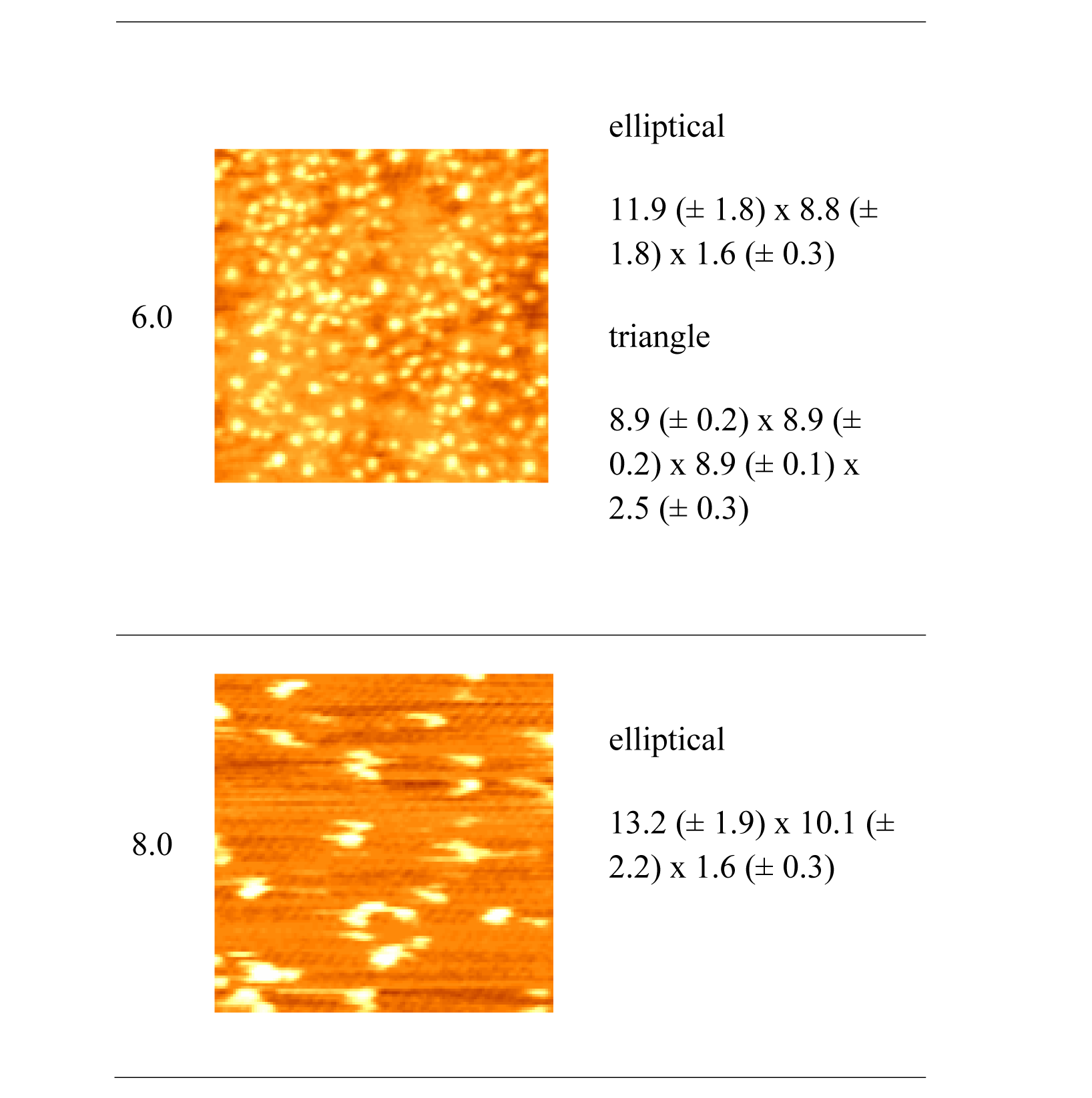
The topography of adsorbed BSA molecules was deposited on a gold surface measured in various pH (c = 1 ppm, I = 0.01 M NaCl for pH = 4.0, 6.0, and 8.0). Images were cropped to 1 μm x 1 μm. The size of the structures was obtained using the ImageJ program, *NIH* (MD, USA). The real dimensions of the long and the short axis of 60 in the plane of the substrate were obtained based on equation 1.

At pH = 4.0, elongated BSA molecules corresponding to the E-form were recorded. In bulk solution, they occur at a pH lower than 3.5. An increase in pH to 6.0 is associated with forming structures of various shapes − those resembling equilateral triangles with dimensions similar to the N-form and elliptical ones. A further increase in pH results in clusters of molecules with a larger cross-sectional area and higher than molecules at pH = 6.0, which means the formation of aggregates. Studies conducted using QCM-D and infrared (IR) spectroscopy reported in the literature have shown that the interaction of BSA with the silicon surface causes changes in protein structure. More minor differences were recorded for BSA containing fatty acids and greater for BSA devoid of these acids, in the case of which, due to weaker side interactions between molecules, more densely packed layers were formed [55]. Research conducted on the adsorption of BSA on silicon (IV) oxide using AFM showed that at pH = 4.0, the particles had a shorter length than those obtained in this work. At pH = 6.0, only triangular-shaped particles corresponding to the natural structure of BSA were obtained, and at pH = 9.0, aggregates [29]. Differences in the structure of molecules adsorbed on silicon (IV) oxide, and the gold surface may be caused by the hydrophilic nature of the silicon (IV) oxide surface, which, compared to the more hydrophobic gold, has a less destabilizing effect on the protein structure [56]. Changes in the shape and dimensions of the adsorbed particles obtained under different pH conditions indicate that interaction with the surface is a factor that induces changes in the structure of BSA, which may be influenced by the properties of the surface itself, such as the degree of hydrophobicity.

### Adsorption Effectiveness and Orientation of BSA Molecules on the Gold Surface Dependent on pH

A change in the charge distribution on the surface of the heterogeneously charged protein, and thus its dipole moment, may be associated with a change in the molecule’s orientation on the surface due to the interaction of other protein regions with the adsorption surface. Therefore, the homogeneity of the charge distribution on the BSA surface depending on the pH was determined by calculating the dipole moment of the BSA molecule using the algorithm proposed by Axel Kohlmeyer [57]. The BSA structure with the identifier 3V03 from the RCSB Protein Data Bank was used. Dipole moment vectors were graphically presented using the VMD program (Supplementary Material Figure SM1). The results of the BSA dipole moment show that at the transition from pH = 3.5 to pH = 9.0, there is a significant change in its value and direction (Figure SM1a). The dipole moment value determined as a simulation time function is highest at pH = 3.5 and is characterized by a 180-degree inverted protein orientation relative to the other states. No significant differences were observed between pH = 6.0 and pH =9.0, either in the molecule’s orientation or the dipole moment values at a given simulation time (Figure SM1b). The information indicates that the molecule’s orientation on the surface can change. To determine the orientation of BSA molecules on the surface and structure of adsorbed films (mono- / multilayers) dependent on pH, the mass of theoretical BSA monolayers () calculated using the random sequential adsorption model (RSA) were compared with the experimental results of the BSA adsorbed mass (). The mass of theoretical BSA monolayers was calculated considering the dimensions of the adsorbed molecules measured by AFM in pH = 4.0 or pH = 6.0. Two main orientations were adopted for BSA: flat-on (when the molecule has the maximum contact area with the gold surface) and side-on (when the molecule adsorbs in such a way as to minimize contact with the gold surface) [58]. The equation used to determine the mass of the BSA monolayer () according to the RSA model is presented below:

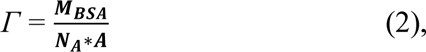

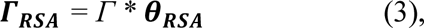

where *Γ* – the mass of BSA monolayer when the surface coverage is equal to 100%, – BSA molar mass equal to 66,430 Da, – Avogadro’s number, *A* – cross-sectional area of adsorbed molecule calculated based on AFM results, and – maximum surface coverage.

The value of for structures with rectangular and triangular cross-sections equal to 0.5, and elliptical structures, it depends on the ratio of the length of the short and long semi-axes according to the concept proposed by J. D. Sherwood [59][60]. The parameters of the molecules combined with the theoretical mass of adsorbed monolayers for protein molecules with selected shapes and orientations are listed in Table SM1. The adsorption of BSA on a gold surface was determined using multi-parameter surface plasmon resonance (MP-SPR). The method allows for determining the adsorption effectiveness of biological samples on metal surfaces in real-time measurements [61]. BSA adsorption was carried out in a pH = 4.0–9.0 with BSA concentration equal to c = 5 ppm in both I = 0.01 M NaCl and I = 0.15 M NaCl, the measurement error was estimated using the total differential method where *Δ(ΔΘ) =* 0.001, and = 0.003 (Figure 2). For the measurements at I = 0.01 M NaCl, the mass of adsorbed BSA in selected pH (pH = 4.0−9.0) is in the range = 30−175 ng/cm^2^. The maximum value of adsorbed mass () is recorded close to BSA isoelectric point at pH = 5.1, which was previously noticed in analogous experiments [27][62][63]. The greater mass of the adsorbed BSA in the isoelectric point range is caused by minimizing the side interactions between the particles and the compact structure of the BSA molecule, which allows more particles to adsorb to the gold surface. Under conditions where both the BSA molecule and the gold surface are negatively charged (above pH > 5.1), BSA adsorption is also observed. This results from electrostatic interactions between the protein domains with the opposite charge in relation to the gold surface and the presence of hydrophobic interactions. The same efficiency of BSA adsorption depending on pH was observed for several other surfaces, such as silicon (IV) oxide, epigallocatechin gallate (EGCG), or glass coated with vinyl alcohol [43–45]. A different trend was observed for measurements at higher ionic strength (I = 0.15 M NaCl). The range of adsorbed mass is narrower (= 88−108 ng/cm^2^), and maximum effectiveness of adsorption was observed at the lowest selected pH equal to pH = 4.0. With increasing pH, a decrease in BSA adsorption efficiency is observed. The change in adsorption efficiency at higher ionic strength is due to charge screening on both the protein and gold surfaces. Both electrostatic and hydrophobic interactions are responsible for BSA adsorption on the gold surface. Adsorption under conditions where both BSA and the surface are negatively charged is possible due to the presence of positively charged lysines (Lys) and arginines (Arg) and negatively charged residues of aspartic (Asp) and glutamic acid (Glu) in the protein structure [64]. Comparison between the mass of theoretical BSA monolayers and the MP-SPR experimental results is shown in Figure 2. Based on the shape of the molecules at the selected pH, it can be concluded that for I = 0.01 M NaCl at pH = 4.0, a film composed of elongated molecules was formed (E-form). Hence, a complete monolayer was not achieved under these conditions. At pH = 5.5, a monolayer of N-form molecules in the flat-on orientation was obtained. At higher pH, both the protein and the gold surface are negatively charged, which results in lower adsorption efficiency and the formation of an incomplete monolayer.

**Figure 2.**
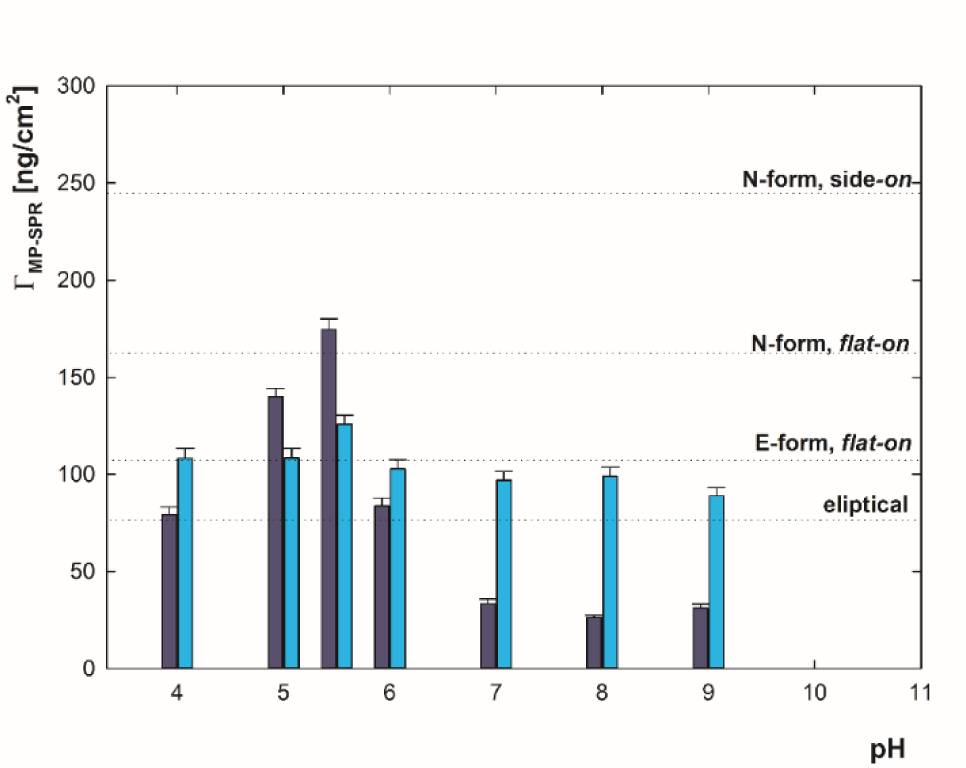
The efficiency of BSA adsorption on the gold surface determined by MP-SPR for pH = 4.0–9.0, c = 5 ppm, I = 0.01 M NaCl (dark blue) or I = 0.15 M NaCl (cyan), the dotted lines represent the mass of theoretical BSA monolayers for selected orientation of BSA molecules calculated based on RSA model.

### Changes in BSS Secondary Structure as a Result of Interactions with the Gold Surface

Measurements of the secondary structure of BSA in solution showed changes in the content of its components depending on pH associated with the opening of BSA structure below pH = 4.0 with a simultaneous decrease of α-helices and increase of β-sheet content (Figure 1b). Additionally, different shapes of BSA molecules adsorbed on gold surface dependent on pH conditions (pH = 4.0, pH = 6.0, pH = 8.0) were confirmed by AFM measurements. Therefore, it was decided to check whether interactions between BSA and the gold surface affect the content of the secondary structure components and what is the direction of these changes. Secondary structure measurements for BSA on the gold surface were performed using Fourier Transform Infrared (FTIR) spectroscopy. Adsorption was carried out on glass plates coated with a gold layer of d = 100 nm. A solution of BSA with a concentration of c = 5 ppm and an ionic strength of I = 0.01M NaCl was adsorbed for 90 minutes in the pH range = 2.0–12.0. Then, it was washed for 90 minutes with I = 0.01 M NaCl, subsequently for 90 minutes in water at the chosen pH. After drying the samples, FTIR spectra were recorded. The position of the bands characteristic for the individual components of the secondary structure was determined based on the location of minima of the II-derivative of the spectra in the range of amide I, which were respectively assigned as 1626 cm^−1^ – β-sheets, 1640 cm^−1^ – disordered structures, 1659 cm^−1^ – α-helices, 1666 cm^−1^ – 3_10_-helices, 1684 cm^−1^ – β-turns and 1694 cm^−1^ – dehydrated β-turns (Figure SM2). The spectra were deconvoluted by fitting the Gauss-Lorentz function at selected positions. The percentage content of a given component of the BSA secondary structure was calculated using the following equation:

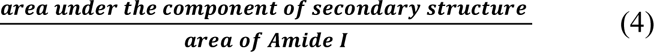

The measurement error was estimated by the measurement uncertainty, which was 5%. The obtained values depending on the pH, are shown in Figure 3b.

**Figure 3.**
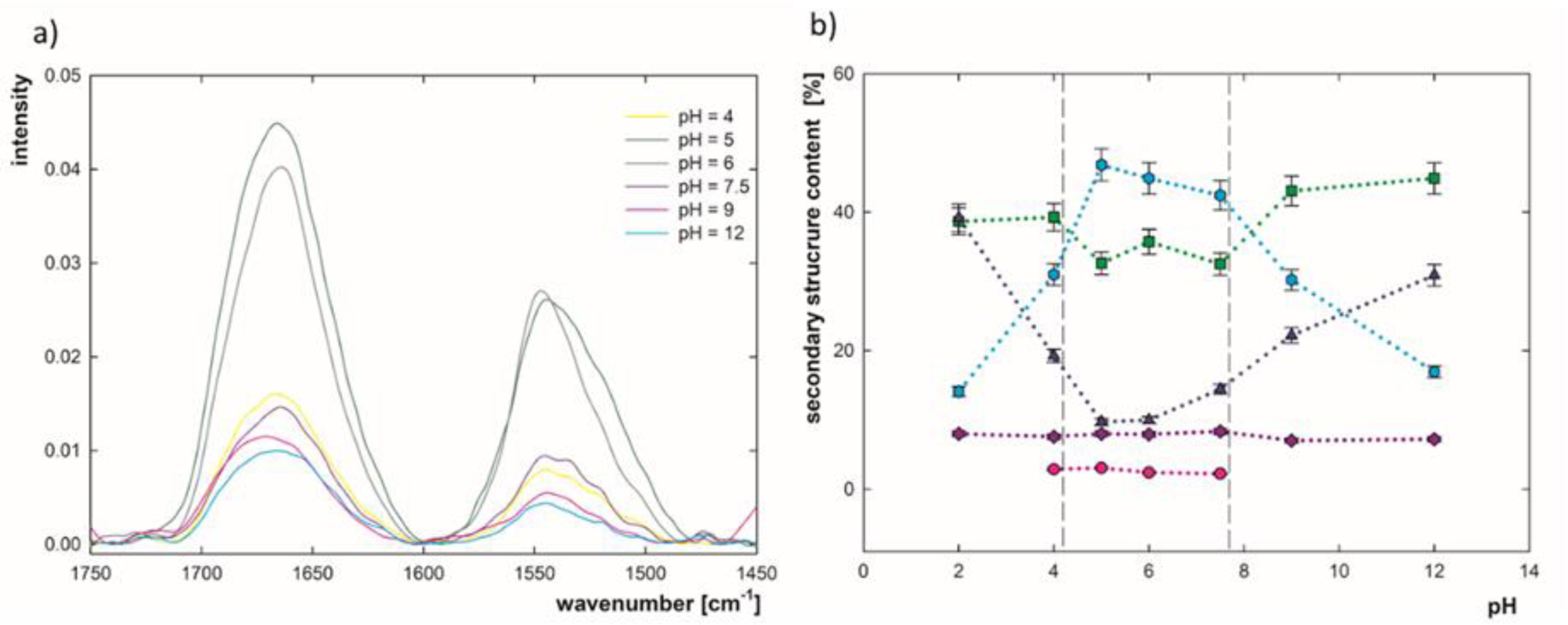
a) FTIR spectra in the wavenumber range of 1450–1750 cm^−1^ for BSA adsorbed on the gold surface depending on pH (c = 5 ppm, I = 0.01 M), b) the percentage content of individual components of the BSA secondary structure in the adsorbed state depending on pH (c = 5 ppm, I = 0.01 M): α-helices (cyan, ⬢), β-turns (green, ▪), disordered structures (navy blue, ▲), β-sheets (purple, ◆), 3_10_-helices (pink, ●).

The strongest changes in the content of secondary structure components for adsorbed BSA depending on pH were observed for α-helices and disordered structures. The α-helices constitute 45% at pH = 5.0–7.5, while at the extreme pH = 2.0 or pH = 12.0 their content drops to 15%. This is related to the increase in the proportion of disordered structures from 10% at the isoelectric point (pH ≈ 5.1) to 40% at pH = 4.0 and 31% at pH = 12.0. In the pH range between 4.0 and 8.0, a lower content of β-turns is observed and, under the same conditions, the presence of 3_10_-helices was also recorded at about 3%. β-sheets are present at a constant level of 8% over the entire pH range, which indicates their high stability. A comparison of the obtained values with the results of the BSA secondary structure in bulk solution shows that the interaction with the gold surface strongly influences the conformation of the protein. Close to the isoelectric point, the content of α-helices remains unchanged after adsorption. A decrease in pH below the isoelectric point is associated with a decrease of α-helices content which occurs both in the bulk solution and in the adsorbed state. However, due to interactions with the surface, the content of helices is lower by almost 10% at pH = 2.0 . However, above pH = 7.0 a greater loss of α-helices after adsorption was observed (around 20% in the highest selected pH). At pH = 2.0 before adsorption, the content of β-sheets is approximately 30% higher. It was also noted that the interaction between BSA and gold resulted in a 30% increase in the content of β-turns and changes in disordered structures at both low and neutral pH. The content of unordered structures on the surface at low pH is higher than in solution, while in the conditions of an isoelectric point lower than in solution. The high content of unordered structures indicates the destabilization of the molecules due to interaction with the gold surface, but the β-sheets remain at a low level, meaning that no BSA aggregation occurred. The increased content of β-turns may prevent aggregation by inhibiting the interactions responsible for exposing protein fragments susceptible to intense conformational changes toward forming aggregates [65]. An analogous direction of changes due to the interaction of BSA with the surface was observed in experiments conducted at pH = 7.4 for BSA adsorbed on both hydrophilic and hydrophobic surfaces [66]. At the same time, conformational changes on hydrophilic surfaces were much smaller. For experiments in which titanium (IV) oxide was used, and measurements were made depending on pH, a decrease in the content of α-helices was recorded, mainly in favor of an increase in the content of disordered structures or β-sheets, intensifying with a decrease in pH. On silicon (IV) oxide, when the pH changed from 7.5 to 2.0, the share of α-helices decreased in favor of β-sheets and β-turns [67][68]. A different direction of changes than for the gold surface may be associated with a different orientation of the BSA molecules and a smaller contact surface of the molecules with hydrophilic surfaces [69]. It was also indicated that the molecules that maintain the triangular shape on the gold surface have a more stable structure than the elliptical ones, for which more significant structural changes are observed, which is confirmed by the comparison of the results obtained in this work at pH = 5.0 and pH = 8.0 [70]. Research on the interaction of BSA with gold nanoparticles in PBS at pH = 7.2 showed that upon protein adsorption, the content of α-helices decreased to 23.5%, while the total content of β-sheets increased to 38% [71]. A different direction of changes may result from differences in data analysis . In the above work, wide ranges of wavenumbers were assigned to individual components of the secondary structure. The presence of β-turns was not considered, and the band corresponding to them was assigned to β-sheets. The interactions of BSA with the surface cause strong changes in the conformation of the protein, the direction of changes and intensity of which depends on the substrate’s wettability and the environment’s pH. In strongly acidic (pH = 2.0–4.0) and strongly alkaline (pH = 9.0–12.0) conditions, intense changes in the secondary structure of BSA on the gold surface are observed, which are also visible by changing the shape of the molecule observed in AFM measurements.

### Confirmation of Structural Changes and Determining the Domain Responsible Through MD Simulations

A comparison of the content of BSA secondary structure components in the adsorbed state with the results of the BSA structure in bulk solution highlights that the interaction with the gold surface strongly influences the conformation of the BSA. Therefore, it was decided to investigate which parts of the protein molecule are most susceptible to such alterations and whether these BSA fragments may be responsible for protein adsorption to the gold surface. For this purpose, molecular dynamics simulations for BSA in different environmental pH conditions were performed. Earlier studies indicate that the flexible segments of the BSA chain participate in the association with the gold surface, the adsorption process is driven by conformational changes, and the internal remodeling of the protein chain enhances interactions with gold through cysteines and aromatic residues [72]. The simulation results show stronger chain vibration under acidic conditions. Intensive fluctuations of the domains IA, IB, IIIA and IIIB were recorded at pH = 3.5 compared to higher pH. For HSA interacting with gold nanorods (AuNR), the correlation of experimental methods (SERS, fluorescence quenching) with predictors of flexible regions (IDP/IDR) was demonstrated, and greater susceptibility to gold interaction of highly flexible segments of human albumin was previously presented [72]. The temperature factors (B-factor) were marked in the secondary structure of the BSA model (PDB: 3V03) with an emphasis on the location of the sulfur atoms (yellow balls, Figure 4a). The parameter allowed visualization of the strongest chain vibrations. The red scale of the upper limit of chain vibration has been selected to highlight the greatest differences in vibration between simulated models (Figure 4a). The strongest vibrations were highlighted graphically based on mean square plots of protein carbon chain fluctuation (RMSF). The maxima of the RMSF plots correspond to the B-factor values, reflecting the relative differences between the structures in the given protonation states. Protein vibrations at pH = 3.5 dominate compared to vibrations occurring at pH = 6.0 and pH = 9.0 (orange curve, Figure 4b). From the mean structures obtained from the RMSF function, the cysteine depth was compared to the vibration of cysteine residues with the greatest depth change during the N-F transition and was marked by “*” (Figure 5c). At pH 3.5, the new cysteines with the greatest transfer to the surface are located in the most charged part of the protein. These regions include the subdomains IA(+8e), IB(+10e) and IIA (+17e) (PDB2PQR Server, empirical pKa predictor for PDB: 3V03). In addition, IA, IB, and IIA subdomains are characterized by the largest number of cysteines that are the source of sulfur groups [73]. Compared to pH 6.0 and 9.0, most of these cysteines (e.g., Cys-53 and Cys-62) lie in regions with very high chain fluctuation (red segments, Figure 4a). Significant increases in subdomain fluctuations, and a large local concentration of disulfide bridges, including the transfer of cysteines towards the surface, make the IA IB and IIA subdomains a highly probable region of contact with the gold surface.

**Figure 4.**
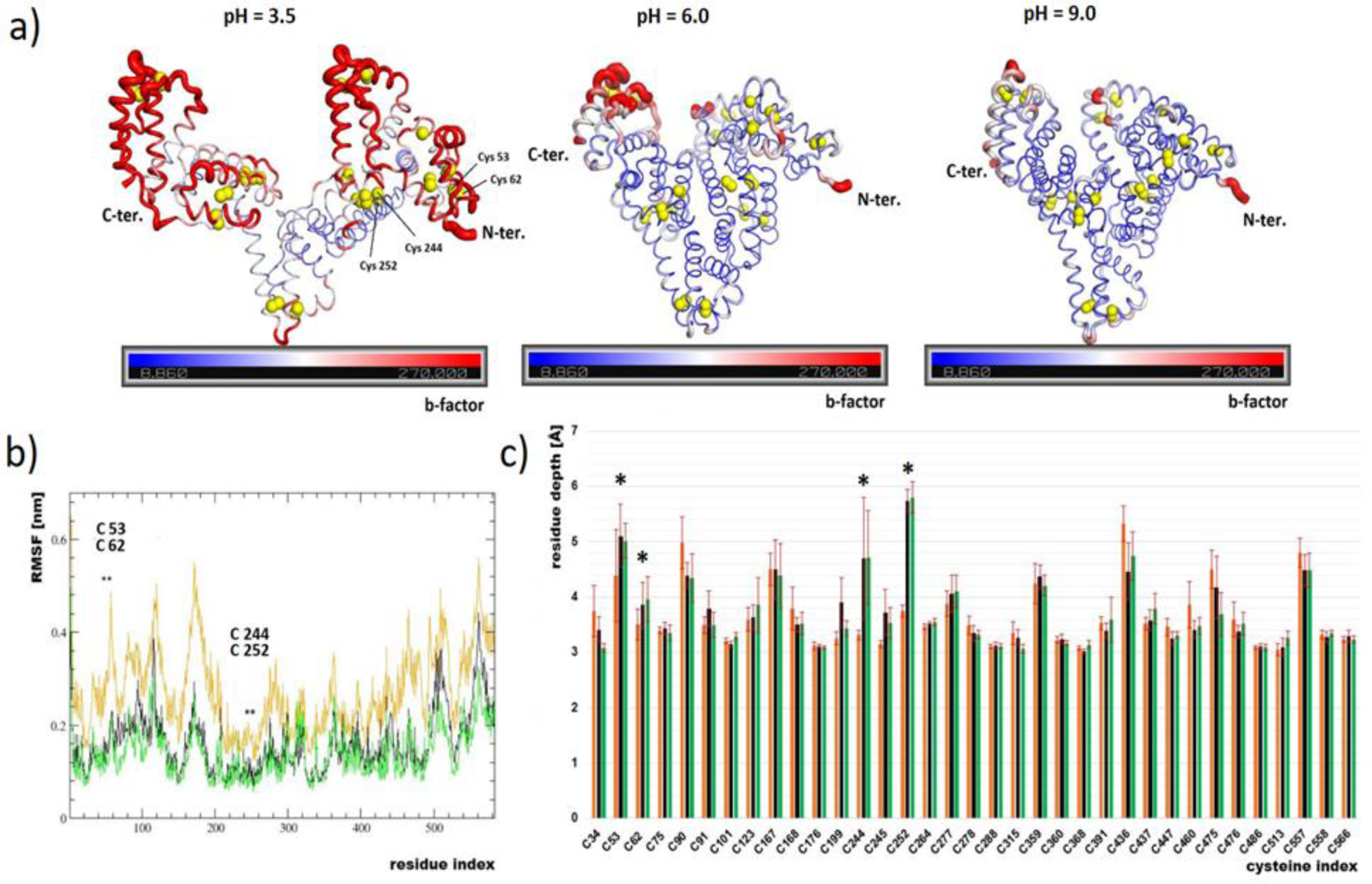
a) B-factor, comparison of vibrations after 200ns of BSA simulations at varying pH levels. Highlighting the distribution of sulphur atoms (yellow balls), cysteine residues with the most significant depth change during the N-F transition, and the final frame of the simulations, b) mean square fluctuations (RMSF), a compilation of vibrations in different protonation states, presentation of vibrations along the chain sequence, *vibration of cysteine residues with the greatest depth change during the N-F transition, (BSA at pH = 3.5 – orange, pH = 6.0 – black, pH= 9.0 – green), c) depth of cysteine residues from the accessible surface (ASA) based on the average structures from the simulation, *cysteine residues with the greatest transfer to the surface during the N-F transition, (BSA at pH 3.5 – orange, pH 6.0 – black, pH 9.0 – green).

**Figure 5.**
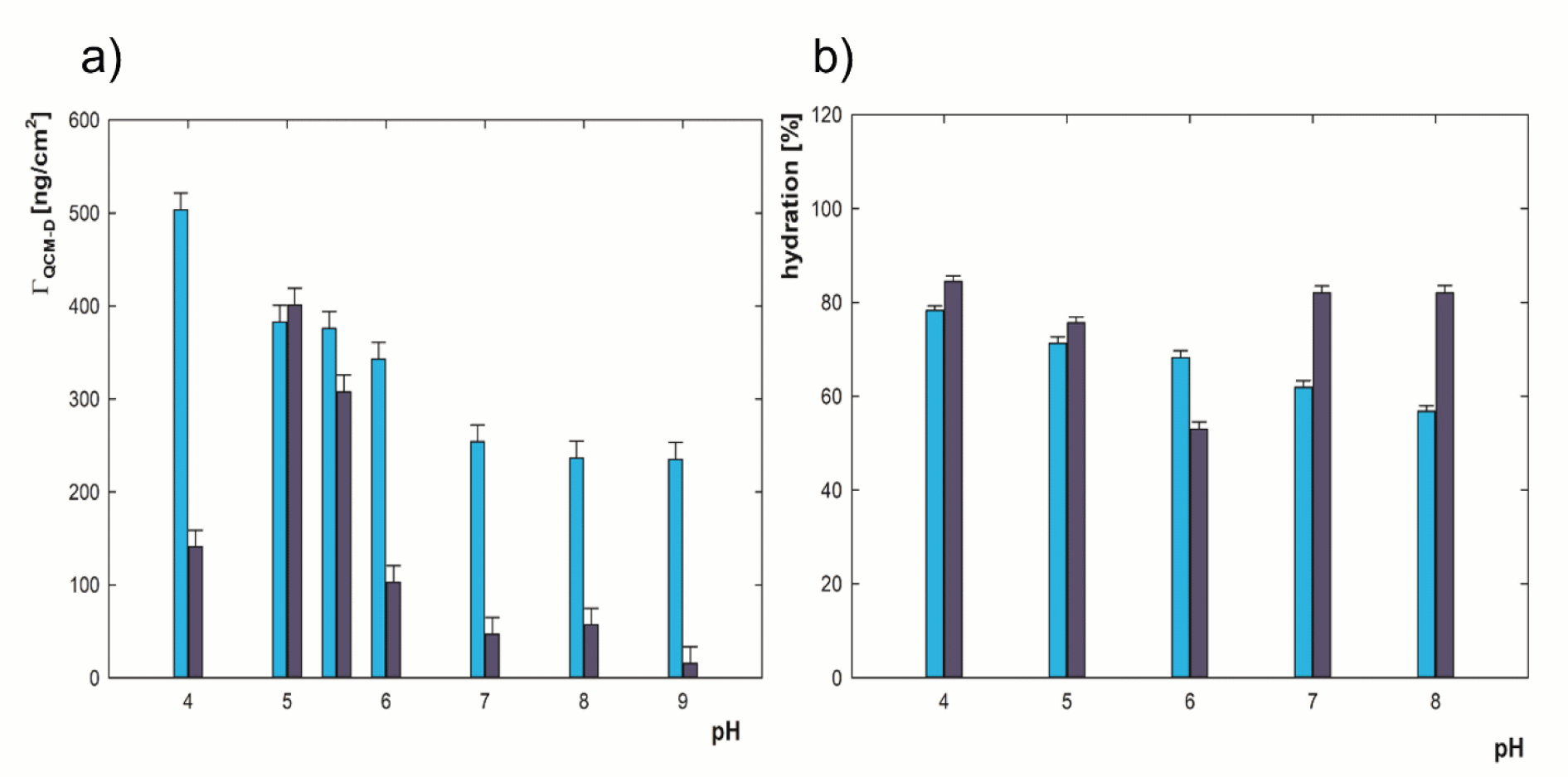
a) The adsorption efficiency of BSA adsorption on the gold surface determined by QCM-D, b) hydration of adsorbed BSA films on the gold surface depending on the pH (for pH = 4.0–9.0, c = 5 ppm, I = 0.01 M NaCl (dark blue) or I = 0.15 M NaCl (cyan)).

Intensive deformations of BSA concern 6 helices covering all protein domains (Figure SM3a). The most deformed helices reach the maximum bending angle above 16° (Figure SM3b), and the remaining helices are below this value of the deformation angle. Specific cysteines are visualized on the helices, characterized by the greatest change in depth towards the molecular surface at the N-F transition, Cys-244 and Cys-252 lie near the intensely deformed region, Cys-53 and Cys-62 lie in the helices the least distorted. Only one free thiol (Cys-34) is available for the protein in N-form on the BSA surface. Usually, inaccessible disulfides can become available for BSA in F- and E-forms to reducing agents or appropriate surfaces [74][75]. Based on the analysis of the Raman spectra of BSA before and after forming AuNPs, the interaction of sulfur groups from the albumin chain with AuNPs was proved. Studies document a significant amplification of Raman vibrations in the presence of gold nanoparticles, which indicates that the appropriate disulfide bridges after adsorption are close to the gold surface, giving a surface-enhanced Raman signal [32][33]. The largest deformations of the secondary structure appear in the centre of the protein (Figure SM3c). The main reason for the increase in BSA chain disorder is electrostatic repulsion, i.e., an increase in the value of the homonymous charge of the opposite domains I and III under acidic conditions. The literature predicts that electrostatic repulsion between the three domains in the protein induces a conformational transition from the N-form to the F-form. It has been calculated that the terminal interdomain spacing of albumin can increase from 3.47±0.12 nm (N-form) to 7.26±0.32 nm (F-form) [51], in these MD simulations, the average distances between domains I and III stabilized at the distance of 5.82±0.68 nm (domain protonation state for pH 3.5 respectively: DI +18e, DII +23e, DIII +20e, simulation temperature 298.15K and ion force 0.01M NaCl, Figure SM3d). The difference in the opening of the structure in relation to previous studies may result from several overlapping factors: ten times higher ionic strengths (0.1M), force field (OPLS/AA), and simulation temperature (300K) [51]. Deformations resulting from the N-F transition were visualized based on the greatest separation of the mass center of domain I from domain III (Figure SM3d).

The availability of new potential thiol groups after the rearrangement of the chain into the F-form can radically change the conformation of the protein on the gold surface, including the architecture of the entire adsorption monolayer. An important factor that can change the surface accessibility of e.g. Cys-244 and Cys-252 is chain deformation, while Cys-53, and Cys-62 can be lifted to the surface due to intense local fluctuations. Greater fluctuation of amino acid residues under acidic conditions led to an intense increase in hydrophobic (brown shades) and hydrophilic (blue shades) surfaces. The results are illustrated in the form of hydrophobicity maps for the last frames of the simulation. Deformations, i.e. stretching and bending of the carbon chain, can be considered as the main cause of the surface increase (Supplementary Materials in Figure SM4a). The dynamic measurement of hydrophobicity indicates that the deformation of the structure at pH =3.5 contributes the greatest increase in both hydrophobic and hydrophilic surface area. No significant differences were observed between pH =6.0 and pH=9.0, with the hydrophilic and hydrophobic surface area values remaining constant over the set simulation time (Figure SM4b).

### The Wettability of Adsorbed BSA Films Dependent on pH

Determination of contact angles for proteins adsorbed on the surface provides information not only regarding the degree of their hydrophilicity/hydrophobicity but can also show deviations in the structure of the obtained protein films. Changes in a protein’s secondary structure or orientation may expose regions that differ in their level of hydrophobicity and may affect surface roughness [76][77]. To check the trend in the level of hydrophobicity for BSA adsorbed on the gold surface as a function of pH (pH range 2.0−12.0) contact angles were measured. The samples were prepared under the same conditions as FTIR measurements. The results are presented in Supplementary Material in Figure SM5. Additionally, the contact angle of the bare gold surface was recorded. The contact angle values were calculated as the average value of 10 measurements for each selected pH, and the measurement error was defined as the standard deviation from the average. A contact angle value of 69.9 ± 1.2° was obtained for a bare gold surface. Similar results for the gold surface were obtained by D. Hong et al. 70.2° [78]. It can be seen from Figure SM5, that there are three characteristic regions: under pH = 4, between pH 4 and 8, and above pH = 8. In pH conditions close to the pH of the native form of BSA in 0.01 M NaCl (pH = 5.5), the values of the contact angles are the highest and reach 59.3–60.2°, which proves that in such conditions, the gold surface with adsorbed BSA is the most hydrophobic. Between pH = 2 and 4, the surface is slightly more hydrophilic. The contact angle drops to 56.9–57.5°. However, from pH = 8 towards higher pH values, there is a strong decrease in the contact angle value to the level of 46° at the highest selected pH. BSA adsorption on the gold surface causes a reduction in the hydrophobicity of the gold surface. The highest adsorption efficiency (amount of adsorbed BSA) occurs in pH = 5.5, so in such conditions the lowest contact angle should be expected, as protein is responsible for increasing surface hydrophilicity. However, the obtained CA values indicate that the trend is reversed. The trend of the contact angle values here is mainly related to changes in BSA structure associated with exposing differential protein regions to the solution under given experimental conditions. It can be seen that the presented changes in selected pH ranges in the graph showing the content of individual components of the secondary structure (Figure 3b) perfectly coincide with the shift in contact angles. Structural changes (formation of an elongated form) that occur at pH = 2.0–4.0 cause the exposure of more hydrophilic parts of the BSA molecule from the surface to the solvent than at pH = 5.0–6.0, what was confirmed by MD simulations. This means that when the molecule is open, the hydrophobic region is responsible for interacting with the surface, and the hydrophilic part is more exposed to the outside in comparison with native pH. Therefore, the formation of aggregates at pH = 8.0 or higher is associated with obtaining even stronger hydrophilic properties. Under such conditions, changes in molecules’ orientation were not observed, therefore, modifications in surface hydrophobicity must be related to secondary structure changes. Similar effects for hydrophobicity were observed for BSA adsorbed on silicon (IV) oxide in different pH conditions [79].

### Hydration of BSA Films Dependent on pH

An essential parameter of the adsorbed protein films is their hydration – the amount of water associated with adsorbed protein molecules due to protein-water interactions. One of the ways to establish the hydration of adsorbed protein films is to perform complementary experiments using the MP-SPR and QCM-D methods. Both methods allow for determining the mass of adsorbed protein on the sensor’s surface. MP-SPR is an optical method sensitive to changes in the refractive index measured at the sensor surface, resulting directly from the adsorption process of molecules. QCM-D is a mechanical (acoustic) method sensitive directly to mass changes on the surface. QCM-D considers not only the mass of adsorbed molecules as MP-SPR ("dry mass"), but also the mass of water present in the film ("wet mass")[80]. Therefore, the comparison of the results obtained with the same adsorption conditions using MP-SPR and QCM-D allows obtaining the hydration of the obtained films (*Γ*_*H*_2_*0*_):

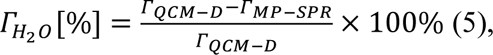

where *Γ*_*MP*−*SPR*_ – the adsorbed mass measured by MP-SPR, *Γ*_*QCM*−*D*_ – adsorbed mass measured by QCM-D, *Γ*_*H*_2_*0*_ – percentage water content [81][82]. The measurement error was calculated using the total differential method, where Δ(Δ*f*) = 1 Hz.

To determine the hydration of the adsorbed films, adsorption efficiency was measured using QCM-D under the same conditions as MP-SPR measurements (c = 5 ppm, pH = 4.0–9.0, I = 0.01 M or I = 0.15 M) and using the same sequence of measurements (10 minutes of baseline, 90 minutes of adsorption, 90 minutes of rinsing by solvent). The QCM-D sensor’s frequency changes can be converted into the adsorbed mass using an appropriate model selected based on the sensor vibration energy dissipation values recorded in the same experiments. When the dissipation exceeds value equal to 10^−6^, it indicates receiving viscoelastic properties by formed layers. The conducted experiments showed that formed BSA films are rigid. In this case, the Sauerbrey model converts the frequency change to adsorbed mass [29][81]. The obtained results are presented in Figure 5a.

It has been proven that for measurements performed at I = 0.01 M NaCl, the maximum efficiency of adsorption occurs at pH = 5, close to the isoelectric point at pH = 5.1 [27]. The mass of adsorbed BSA on the gold surface varies from *Γ*_*QCM*−*D*_= 20 ng/cm^2^ to *Γ*_*QCM*−*D*_ = 400 ng/cm^2^ in the pH range = 4.0–9.0. Under conditions where both the BSA molecule and the gold surface are negatively charged (pH > 5.1), BSA adsorption is still observed. The same effect occurred for MP-SPR measurements and is caused by the heterogeneous charge distribution on the protein surface and the possibility of local hydrophobic interactions. In the QCM-D experiments, at the concentration of BSA equal to 5 ppm, I = 0.01 M NaCl on the hydrophilic surface of silicon (IV) oxide in a wide range of pH, lower adsorption efficiency was recorded (***Γ***_***QCM***−***D***_= 10 ng/cm^2^ at pH = 5.0–6.0) than for the gold surface. Hydrophobic surfaces enhance the association of protein molecules on the surface by forming hydrogen bonds between the BSA and the surface. For higher ionic strength (I = 0.15 M NaCl), the range of mass values of adsorbed BSA films is in the range from ***Γ***_***QCM***−***D***_= 230 ng/cm^2^ to ***Γ***_***QCM***−***D***_= 500 ng/cm^2^. With increasing pH, a decrease in BSA adsorption efficiency is observed. Hydration was calculated by comparing MP-SPR results presented in Figure 2 and QCM-D in Figure 5a. The tendency of BSA adsorption efficiency for I = 0.01 M NaCl obtained using both methods is similar. The adsorption maximum occurs in the range of the isoelectric point. The decrease in the adsorbed BSA mass occurs while moving away from the zero point of the BSA charge. For I = 0.01 M NaCl, between the lowest and the highest measured value of adsorbed mass, an eleven- and seven-fold increase was recorded, respectively, for the results obtained with QCM-D and MP-SPR, respectively, which indicates differences in films; hydration depending on pH. In the case of the measurement series for the ionic strength I = 0.15 M NaCl, the recorded mass values depending on the pH of the solution for the QCM-D and MP-SPR methods have a different path. The increase in ionic strength significantly affects the degree of hydration of the BSA film. Based on equation 7, the percentage of water content in the adsorbed BSA films on the gold surface, depending on pH was calculated. For I = 0.01 M NaCl, hydration changes in the range *Γ*_*H*_2_*0*_ = 48–85%, they are obtaining minimum values at pH = 5.5, i.e., in conditions close to the BSA isoelectric point. In such conditions, the highest degree of surface coverage was recorded. Under conditions of lower and higher pH, hydration increases rapidly. At pH = 4.0, where *θ* = 77%, hydration reaches *Γ*_*H*_2_*0*_ = 85%, while at pH = 8.0, where coverage is *θ* = 17% water content is *Γ*_*H*_2_*0*_ = 82 %. For the ionic strength I = 0.15 M NaCl at pH = 4.0, where a complete flat-on monolayer of molecules in E-form was obtained (*Γ*_*MP*−*SPR*_ = 112 ng/cm^2^) with the highest water content observed. Despite similar coverage for pH = 6.0 and pH = 8.0, there are structural variations in the formed films due to changes in the shape of the molecules and changes in the protonation of BSA. The differences in structure result in a decrease in hydration level to *Γ*_*H*_2_*0*_ = 67% and *Γ*_*H*_2_*0*_ = 57% at pH = 6.0 and pH = 8.0, respectively. Increasing the ionic strength (I = 0.15 M NaCl) has a stabilizing effect on the protein films, preventing changes in the structure and the viscoelastic properties [86][87]. For both ionic strengths, maximum hydration occurs at pH = 4.0 when molecules with an elongated structure are observed (E-form) in a flat-on orientation. This structure allows more water molecules to bind to the BSA molecule. BSA adsorbed on hydrophilic silicon oxide (IV) hydration is *Γ*_*H*_2_*0*_ = 85–95% of water with adsorbed mass in the range of *Γ*_*MP*−*SPR*_ = 5–75 ng/cm^2^, while on the hydrophobic surface modified with methyl groups, hydration remains at *Γ*_*H*_2_*0*_ = 75% in a wide range of protein films; masses (*Γ*_*MP*−*SPR*_ = 50–175 ng/cm^2^). Presented in the literature, measurements of the change in the vibration frequency of the QCM-D sensor for I = 0.15 M NaCl showed that the water content in the BSA films adsorbed on the gold surface increases 1.4 times when the pH changes from 4.5 to 7.0 and depends on the ionic strength [57]. The results are consistent with the hydration level shown in Figure 5b. The mass content ratio increased 1.25 times from pH = 4.0 to pH = 7.0. Adsorption conditions and the structure of molecules have a stronger effect on the hydration of BSA films than the degree of monolayer filling. The increase in ionic strength (I = 0.15 M NaCl) results in more vital compensation of the surface charge of protein and gold by counterions and, thus smaller changes in hydration depending on pH.

### Adsorption Efficiency, Orientation and Hydration of BSA Molecules on the Surface Depending on the Concentration

In order to determine the adsorption efficiency and orientation of adsorbed BSA molecules on the gold surface depending on the BSA concentration of BSA, the MP-SPR method was used. Measurements were carried out in the c = 5–200 ppm concentration range at pH = 6.0 for I = 0.01 M NaCl (Figure SM6a). The adsorbed mass rises with the increase of the concentration in the range c = 5–75 ppm. However, the film becomes saturated at an even higher concentration of BSA (c = 75–200 ppm). For the concentration c = 5 ppm, after 90 minutes of adsorption and 90 minutes of rinsing with a solution of I = 0.01 M NaCl, the adsorbed mass was equal to Γ_MP−SPR_= 85 ng/cm^2^, whereas the curves representing the highest concentrations (c = 75–200 ppm) remain at range Γ_MP−SPR_= 140–170 ng/cm^2^, what is presented in Figure SM6a. In Figure SM6b, the results are compiled after 90 minutes of BSA adsorption (dark blue bars) and after another 90 minutes of rinsing with solvent (cyan bars). Desorption rises with increasing BSA concentration but remains low and does not exceed 5%, while its average value is 2%, which means that BSA adsorption on the gold surface is irreversible. The horizontal dashed lines correspond to the mass of the BSA monolayer in different orientations, which were previously calculated based on dimensions of adsorbed BSA molecules obtained by the AFM and RSA model (Table SM1). N-form and elliptical structures identified at pH = 6.0 for which the mass of monolayer is 161.1 ng/cm^2^ or 247.4 ng/cm^2^, for N-form molecules in flat-on or side-on orientation, respectively, and 76.8 ng/cm^2^ for elliptical structures. For c = 5 ppm, approximately *θ* = 52% of the maximum surface coverage of the monolayer was obtained for molecules with the N-form in the flat-on orientation. In the case of a ten-fold increase in concentration, the surface coverage was *θ* = 80%. For the concentration equal to c = 100 ppm, the mass Γ_MP−SPR_= 158 ng/cm^2^ was recorded, which proves the formation of a monolayer of N-form molecules in the flat-on orientation. An increase in BSA concentration causes the association of subsequent molecules to the monolayer, but they are desorbed due to rinsing the formed layer with a solvent. The received orientation of BSA molecules adsorbed on a gold surface is consistent with studies on the adsorption of BSA on the surface of polystyrene having hydrophobic properties. Even a significant increase in protein concentration does not change the orientation of molecules and the film structure. In the same research, T. Wangkam and his team, based on film thickness measurements made by AFM in the contact mode, also found that the molecules are oriented in the flat-on position [85]. Studies conducted using molecular dynamics (MD) simulations have shown that for the adsorption of negatively charged BSA to the negatively charged surface, domain III is mainly responsible, which, unlike domains II and III of BSA, is positively charged [86]. The positively charged lysines included in the III domain of BSA (Lys535, Lys537) strongly anchor on the surface and significantly impact the stable adsorption of BSA on the surface. In this case, the negatively charged domains face the solvent [86]. These results are consistent with the conclusions drawn for BSA MD simulations on gold nanorods, according to which the part responsible for BSA adsorption to the surface formed primarily from disulfide bonds present in domain III of BSA [87]. The conducted research shows that BSA particles are irreversibly adsorbed on the gold surface and do not undergo reorganization. These studies confirm the preferred orientation of BSA molecules in relation to negatively charged surfaces. BSA does not form multilayers, proving weak interactions between the formed protein film and BSA molecules associated with the resulting layer.

The hydration of the BSA films adsorbed from different protein concentrations was determined by comparing the results of the adsorbed mass using the MP-SPR and QCM-D methods using equation 7 (Figure 6). For this purpose, similar BSA adsorption experiments were performed depending on the concentration using QCM-D (Figure SM6c and SM6d) to those achieved using MP-SPR. The tendency of the adsorbed mass value measured by QCM-D and MP-SPR is similar. The adsorbed mass remains constant starting from a concentration of c = 100 ppm. The mass values obtained with the QCM-D are more than three times higher than those obtained with the MP-SPR. The films formed at a protein concentration c = 75–200 ppm are characterized by stable hydration levels, the average value of which is Γ_H2O_ = 67%. BSA adsorption in the concentration range of c =75–100ppm resulted in changes in surface coverage by 20 percentage points. However, the variations do not affect the hydration level of the formed protein films on the hydrophobic gold surface, which is consistent with the results reported by M. M. Ouberai et al. [69]. In their work, the hydration values of the obtained films are 10 percentage points higher, which may be due to the nature of the adsorption surface, which was functionalized with methyl groups [69]. The same work shows that with a surface coverage of *θ* = 50%, the amount of water contained in the films is Γ_H2O_ = 74% and is lower by about four percentage points than obtaining a complete monolayer . Still, it is not as big a difference as in the case of the presented measurement for BSA adsorbed on the gold surface, where the difference with the same surface coverage is up to 17% [69]. Moreover, when the monolayer is forming a monolayer in the range of low surface coverage, higher hydration is often observed, which is caused by water binding not only to protein molecules but also to the adsorption surface [69][74][88].

**Figure 6.**
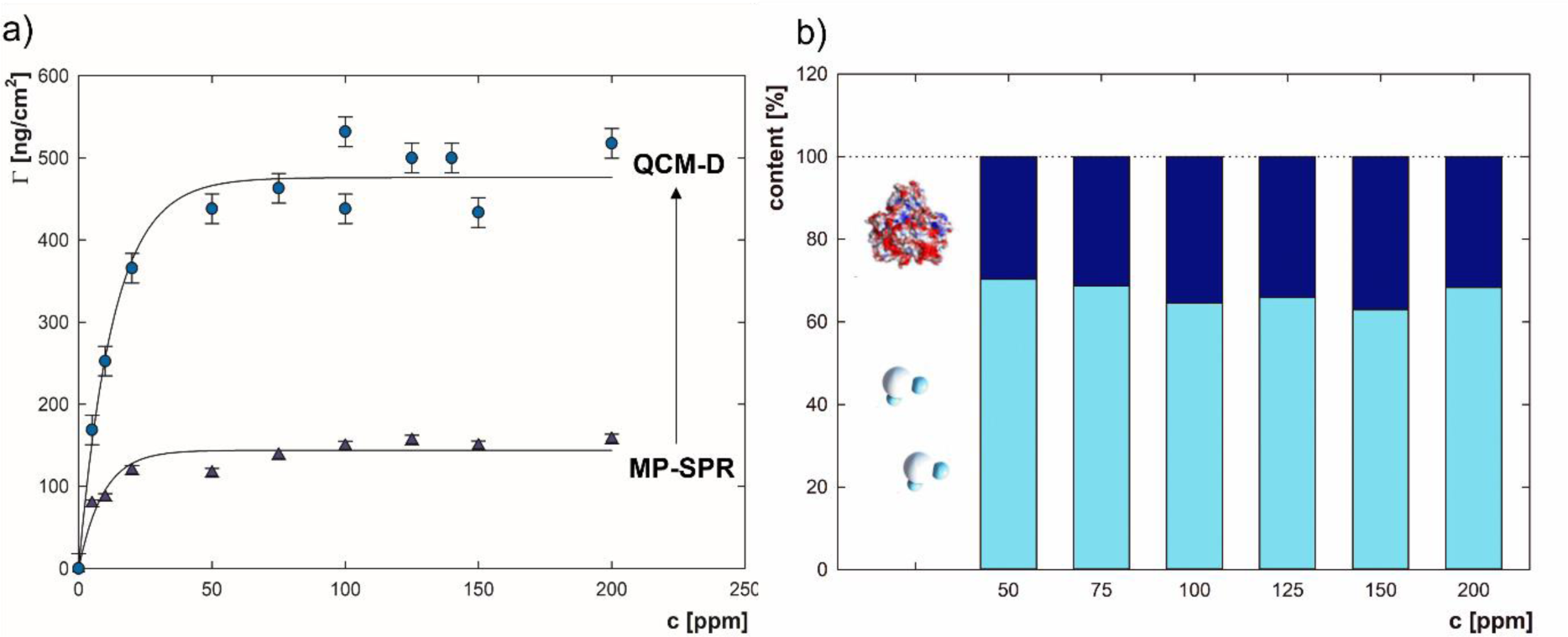
The percentage content of water (blue) and BSA (navy blue) in the mass of adsorbed BSA films dependent on protein concentration.

### Effect of Protein Concentration on the Secondary Structure of Adsorbed BSA

The effect of BSA concentration on the secondary structure of BSA adsorbed on the gold surface was determined using FTIR measurements (c = 500 ppm, I = 0.01 M NaCl, pH = 6.0). The recorded spectra were deconvoluted, and then, the content of particular secondary structure components was calculated using equation 6. Based on the MP-SPR measurements and theoretical calculations, it was established that the complete monolayer is formed starting at c = 100 ppm, and in the entire concentration range, the molecules are in the flat-on orientation. In the range of concentrations higher than c = 50 ppm, the BSA structure on the gold surface was stable and consisted of 56% α-helices, 26% β-turns, 7% – disordered structures, 6% – β-sheets, and 3_10_-helices about 2% (Supplementary Material, Figure SM5). At lower concentrations, slight changes in the α-helices (52% for c = 5 ppm) were observed, favouring a higher content of β-turns. With a higher degree of surface coverage, the secondary structure of BSA is more similar to the native structure of the protein than under conditions of lower surface coverage. In situations with more molecules per unit area, the side interactions between the adsorbed molecules can stabilize the conformation. A similar trend was observed for BSA adsorbed on the hematite surface at pH = 5.0, I = 0.01 M NaCl. A smaller content of α-helices and a higher content of β-structures were confirmed at a lower BSA concentration (c = 30 ppm) than the experiments at a higher concentration (c = 70 ppm). Above this concentration, the band’s intensity from aggregated elements increased, and the side chains were less exposed. Based on the course of the adsorption isotherm, it was found to be associated with forming a multilayer [89]. No multilayers were obtained in the case of the presented research for BSA adsorption on the gold surface. A higher content of α-helices with a higher surface coverage was obtained in the case of BSA adsorbed on hematite particles at pH = 5.7 [90]. The structure of adsorbed BSA molecules depends on the degree of surface coverage and results from the influence of protein-surface interactions as well as protein-protein interactions [91].

## Conclusion

The main aim of the research was to identify the conformational and structural effects of the adsorption of the BSA molecule on the gold surface. The topography of the BSA particles adsorbed on the gold surface obtained by AFM shows that the size and shape of the particles vary with pH. At pH = 4.0, the BSA molecule is in the elongated E-type form, while in solution, this form is observed only for a pH lower than 3.5. At pH = 6.0, the protein has a triangular cross-section typical of the N form. A further increase in pH leads to the formation of aggregated forms. The dimensions of the adsorbed BSA molecule obtained by AFM were used to estimate the theoretical mass of BSA monolayers with different orientations using the RSA model. Comparing the theoretical values with MP-SPR results confirms that the molecules adsorb in a planar orientation, independent of pH and protein concentration. In addition, no multilayer formation was observed. The correlation of changes in the secondary structure in solution (CD) and adsorbed state (FTIR) confirmed that the interaction of BSA with the gold surface causes substantial modifications of protein conformation. The direction and intensity of these changes strongly depend on pH, while the protein concentration does not have such a significant effect. In strongly acidic (pH = 2.0–4.0) and alkaline (pH = 9.0–12.0) conditions, an intensification of changes in the secondary structure of BSA was observed towards a lower content of α-helices and a higher proportion of disordered forms. The increased range of disordered structures indicates the destabilization of molecules due to interaction with the gold surface. At the same time, the content of β-sheets is low, which means no protein aggregation. The increased content of β-turns may prevent aggregation by inhibiting the interactions responsible for exposing protein fragments susceptible to significant conformational changes. The BSA at the molecular level was visualized using MD simulations. MD simulations showed that BSA adsorption at acidic pH involved more sulfur amino acids interacting with the gold surface than at alkaline pH. Disulfide bridges significantly affect the stabilization of the protein structure. Therefore, the participation of cysteines in the adsorption process may result from conformational changes. The above changes are also visible in the values of the contact angle of the gold surface covered with protein, which indicates the level of hydrophobicity of the obtained layers. In addition, the adsorption conditions and molecular structure have a decisive influence on the hydration of BSA layers than the degree of surface coverage.

## Supporting Information

Supplementary Materials file is attached.

## Supporting information

Supplementary Material Bio

## Acknowledgements

Funding: The presented work was supported by Grant NCN PRELUDIUM 2020/37/N/ST4/02132, and NCN OPUS 2021/41/B/ST5/02233. Paulina Komorek acknowledges the support of InterDokMed project no. POWR.03.02.00-00-I013/16. This study was partly supported by PLGrid Core project no PLG/2022/015447.

## Notes

### Competing Interest Statement

The authors have declared no competing interest.

