## Supplementary Material Bio for "Changes in Secondary Structure and Properties of Bovine Serum Albumin as a Result of Interactions with Gold Surface"

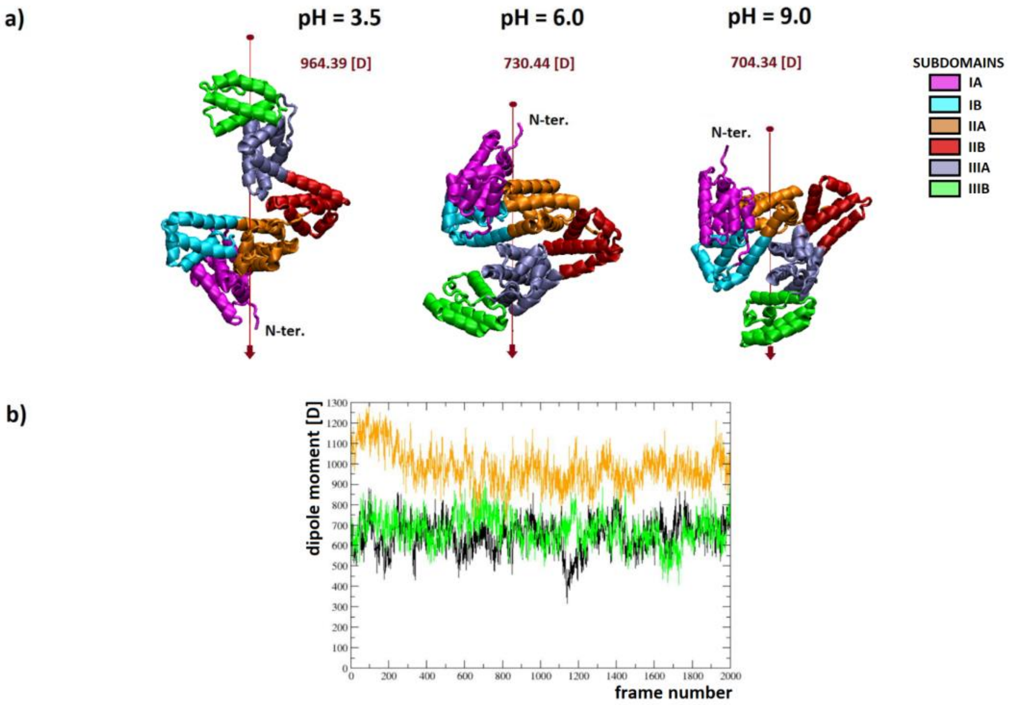


Figure SM1. a) The dipole moment vector of the BSA molecule (last frames) dependent on pH for I = 0.01 M, b) dipole moment value as a function of simulation time (pH = 3.5 – orange, pH = 6.0 – black, pH = 9.0 – green).

Table SM1. Dimensions of adsorbed BSA molecules on the gold surface together with masses of BSA monolayers for individual shapes of molecules and their selected orientations calculated based on the RSA model (*Γ* – the mass of BSA monolayer when the surface coverage is equal to 100%, *A* – cross-sectional area of adsorbed molecule calculated based on molecules dimensions obtained by AFM measurements, and – maximum surface coverage).

measurements, and – maximum surface coverage).

| **Shape of molecule** | **E-form** | | **elliptical** | **N-form** | |
|---|---|---|---|---|---|
| **spatial structure** | 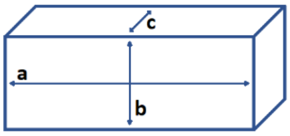 | | 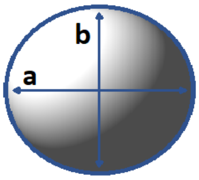 | 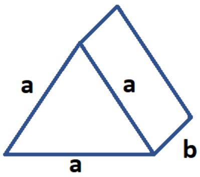 | |
| **cross-sectional dimensions [nm]** | a = 21.2 (± 2.5)<br>b = 2,5 (± 0.3) | b = 2.5 (± 0.3)<br>c = 1.2 (± 0.2) | a = 11.9 (± 1.8)<br>b = 8.8 (± 1.8) | A = 8.9 (± 0,2) | a = 8.9 (± 0.2)<br>b = 2.5 (± 0.3) |
| **orientation** | *flat-on* | *side-on* | | *flat-on* | *side-on* |
| **A [nm$^2$]** | 53.0 ± 12.6 | 3.0 ± 0.9 | 82.1 ± 37 | 34.3 ± 14.2 | 22.3 ± 3.2 |
| **Γ [ng/cm$^2$]** | 208.1 ± 28.2 | 3677 ± 628.8 | 134.8 ± 30.4 | 322.1 ± 76.0 | 494.7 ± 40.5 |
| | 0.5 | 0.5 | 0.57 | 0.5 | 0.5 |
| **$Γ_{RSA}$ [ng/cm$^2$]** | 104.1 ± 14.1 | 1839 ± 314.4 | 76.8 ± 19.7 | 161.1 ± 38.0 | 247.4 ± 20.3 |

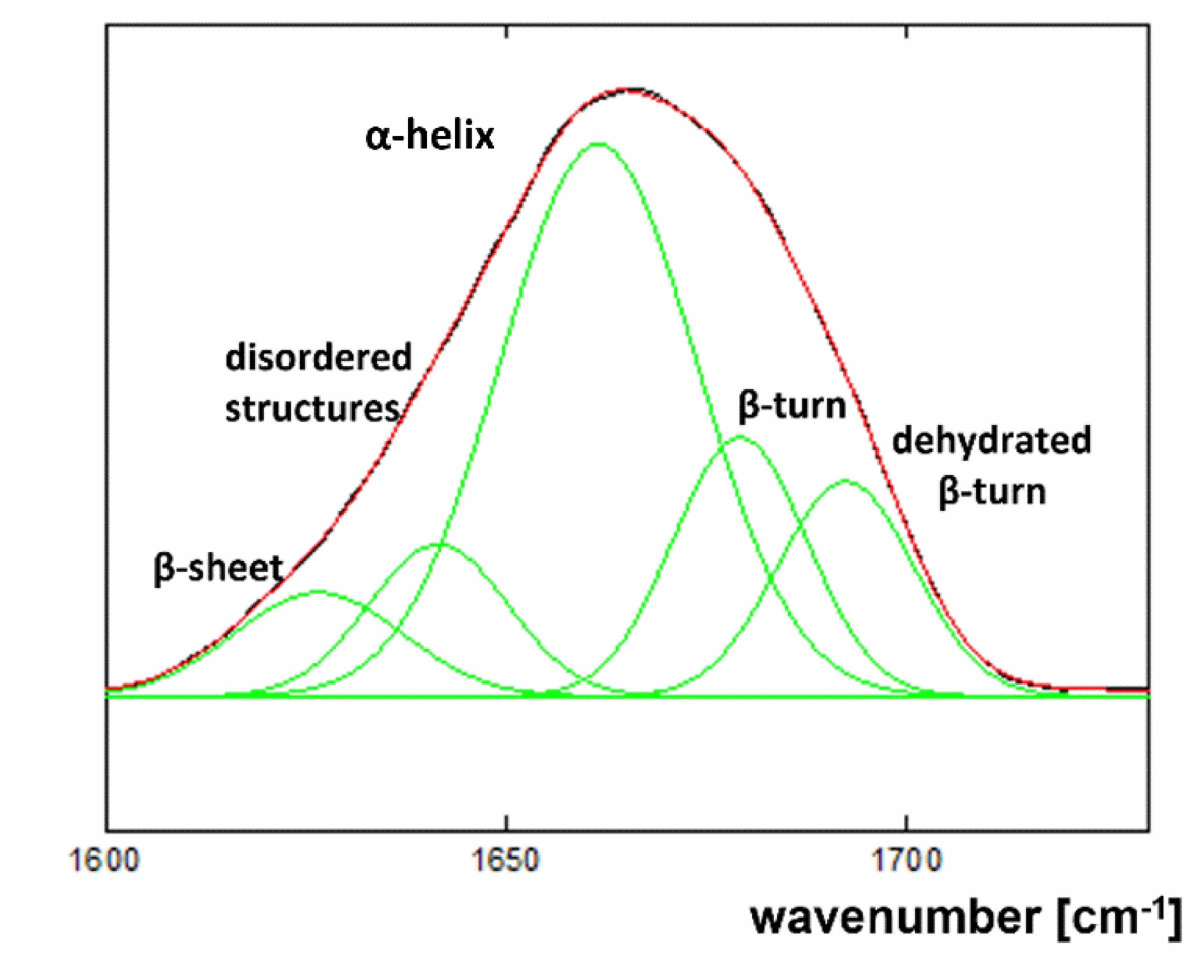


Figure SM2. A scheme of fitted bands corresponding to the elements of the BSA secondary structure to the area of amide I.

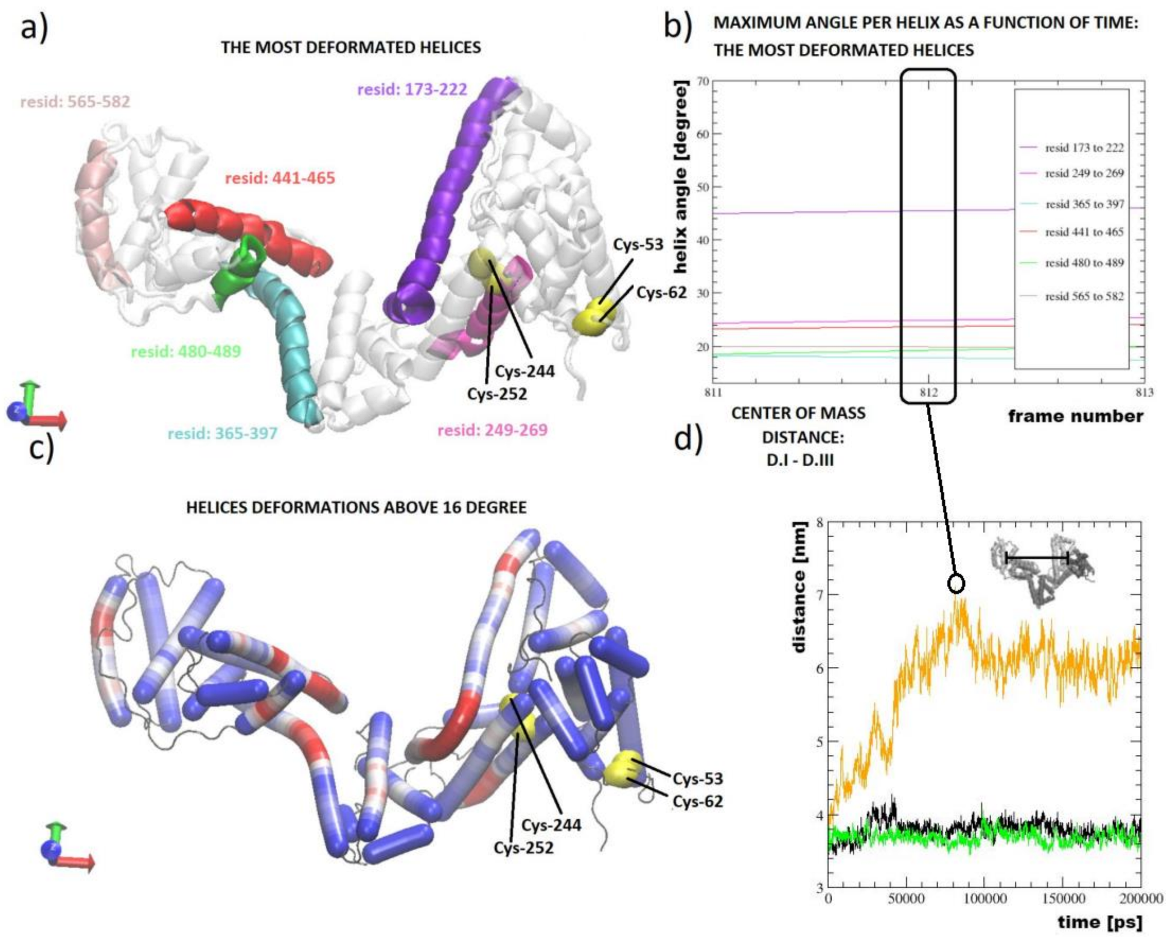


Figure SM3. a) Visualization of the most deformed helices during the N-F structural transition (marked with colors), b) the six most deformed helices reaching maximum bend angles above 16° (the colors of the graph correspond to the graphical visualization on the structure, the black border corresponds to the frame of the greatest flare domains I and III), c) visualization of deformations along the BSA helices, deformations above the maximum threshold of 16° are underlined in red, d) summary of the mean distance between the mass centers of domain I and domain III, visualization of the transition between the N-F form, (BSA at pH = 3.5 – orange, pH = 6.0 – black, pH = 9.0 – green).

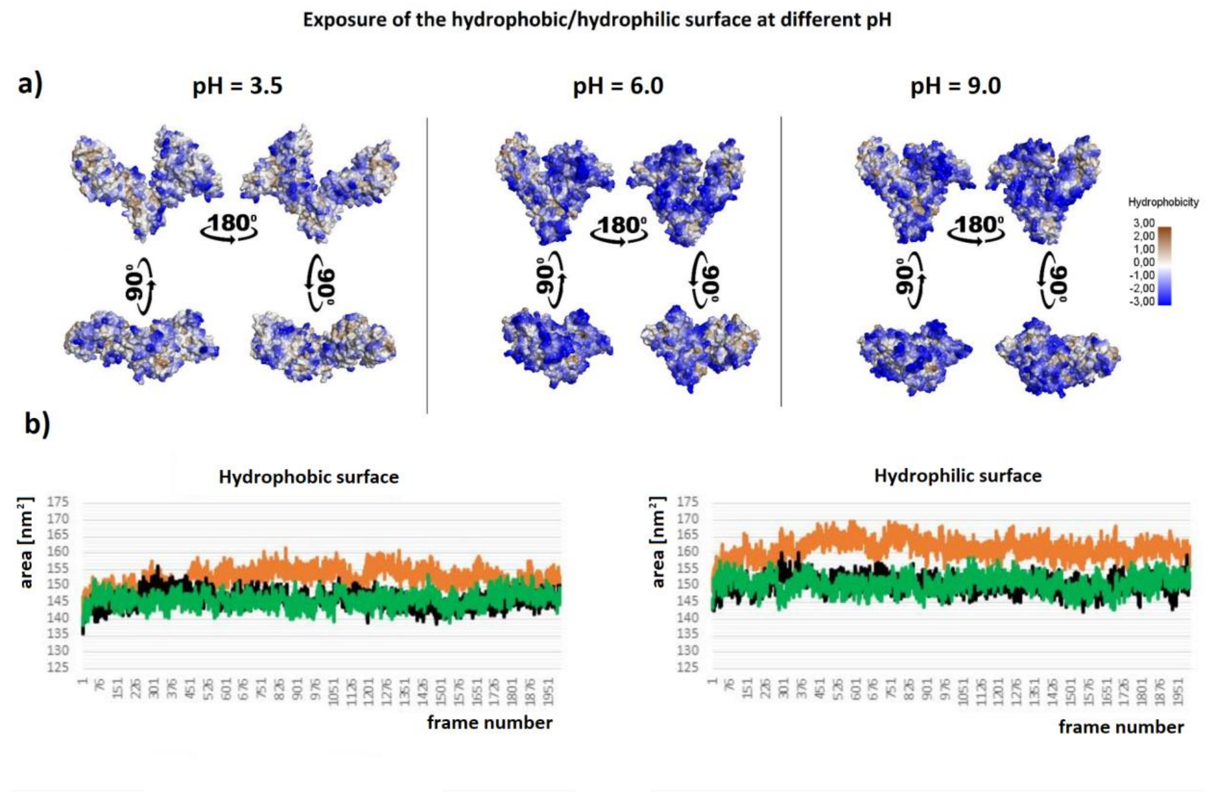


Figure SM4. a) Comparison of BSA hydrophobicity maps in selected protonation states (pH = 3.5, pH = 6.0, pH = 9.0), last frames after 200 ns simulation (I=0.01 M, T=298.15 K, p=1 Bar, forcefield CHARMM36m, water model TIP3P), b) comparison of changes in the hydrophilic and hydrophobic surface area of a protein as a function of simulation time in given protonation states (pH = 3.5 orange, pH = 6.0 black, pH = 9.0 green).

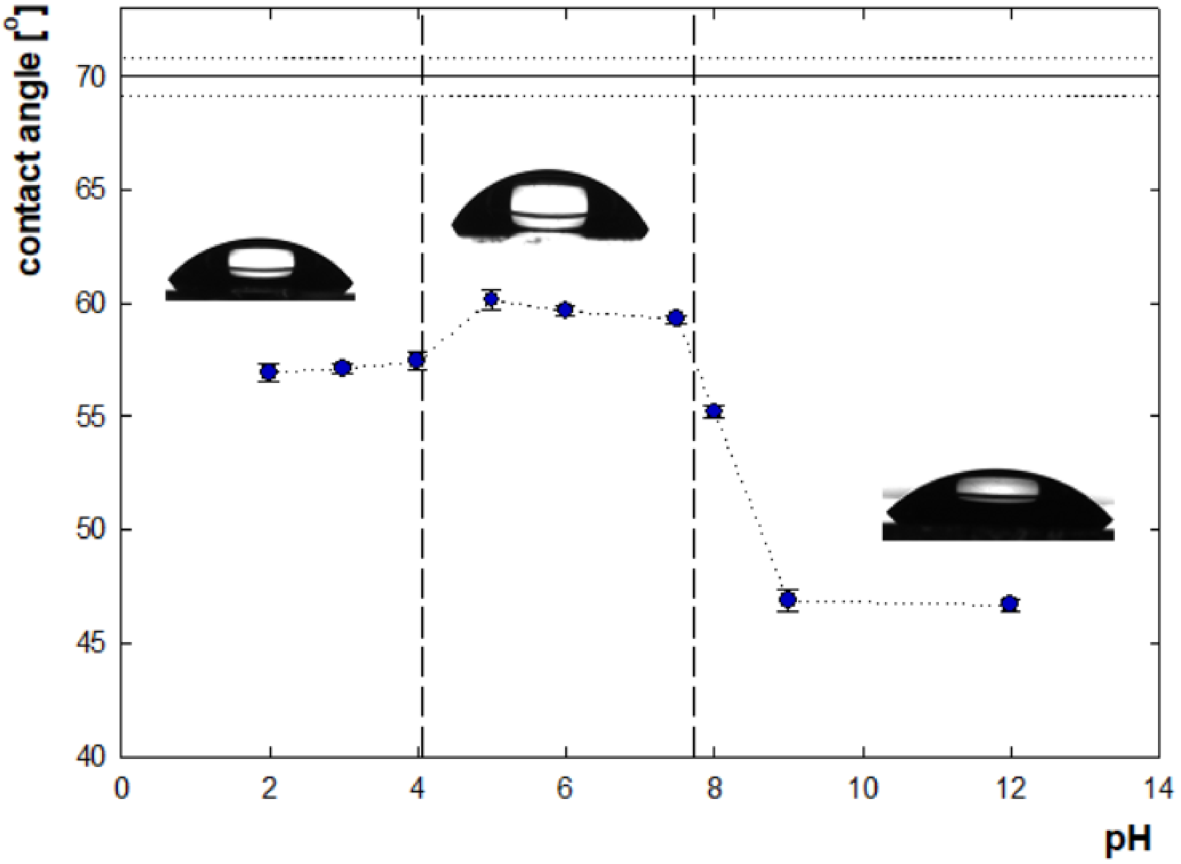


Figure SM5. The contact angle values for BSA adsorbed on gold surface dependent on pH (I = 0.01 M NaCl, c = 5 ppm), the horizontal line indicates the contact angle for the bare gold surface.

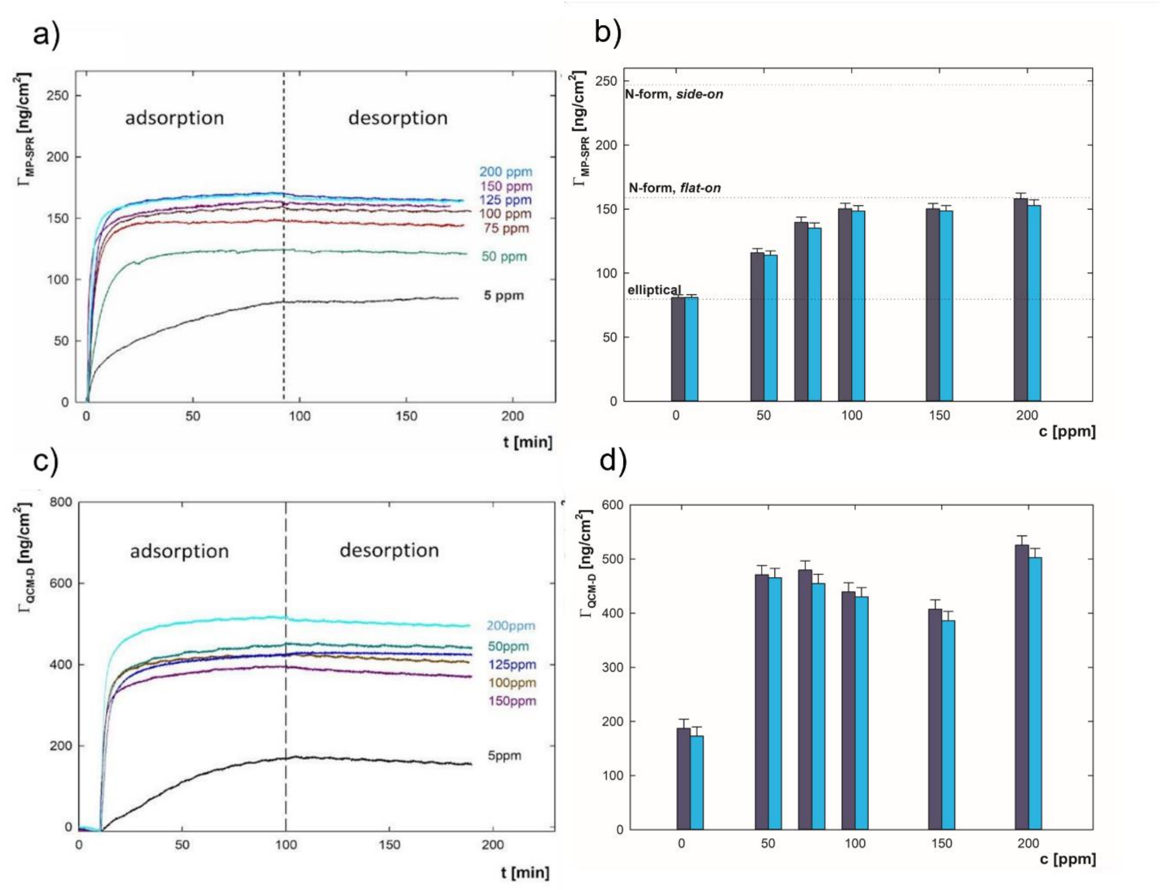


Figure SM6. The efficiency of BSA adsorption measured by MP-SPR and QCM-D a) changes in BSA adsorbed mass (MP-SPR) over time dependent on concentration, b) comparison mass of adsorbed BSA after 90 minutes of adsorption (dark blue) and subsequent 90 minutes of rinsing with solvent (cyan) measured by MP-SPR dependent on pH c) changes in BSA adsorbed mass (QCM-D) over time dependent on concentration. The dotted lines indicate the mass of the BSA monolayers for flat-on or side-on orientation (Table 2), d) comparison mass of adsorbed BSA after 90 minutes of adsorption (dark blue) and subsequent 90 minutes of rinsing with solvent (cyan) measured by QCM-D dependent on pH (pH = 6.0, I = 0.01 M NaCl in the concentration range c = 5–200 ppm).

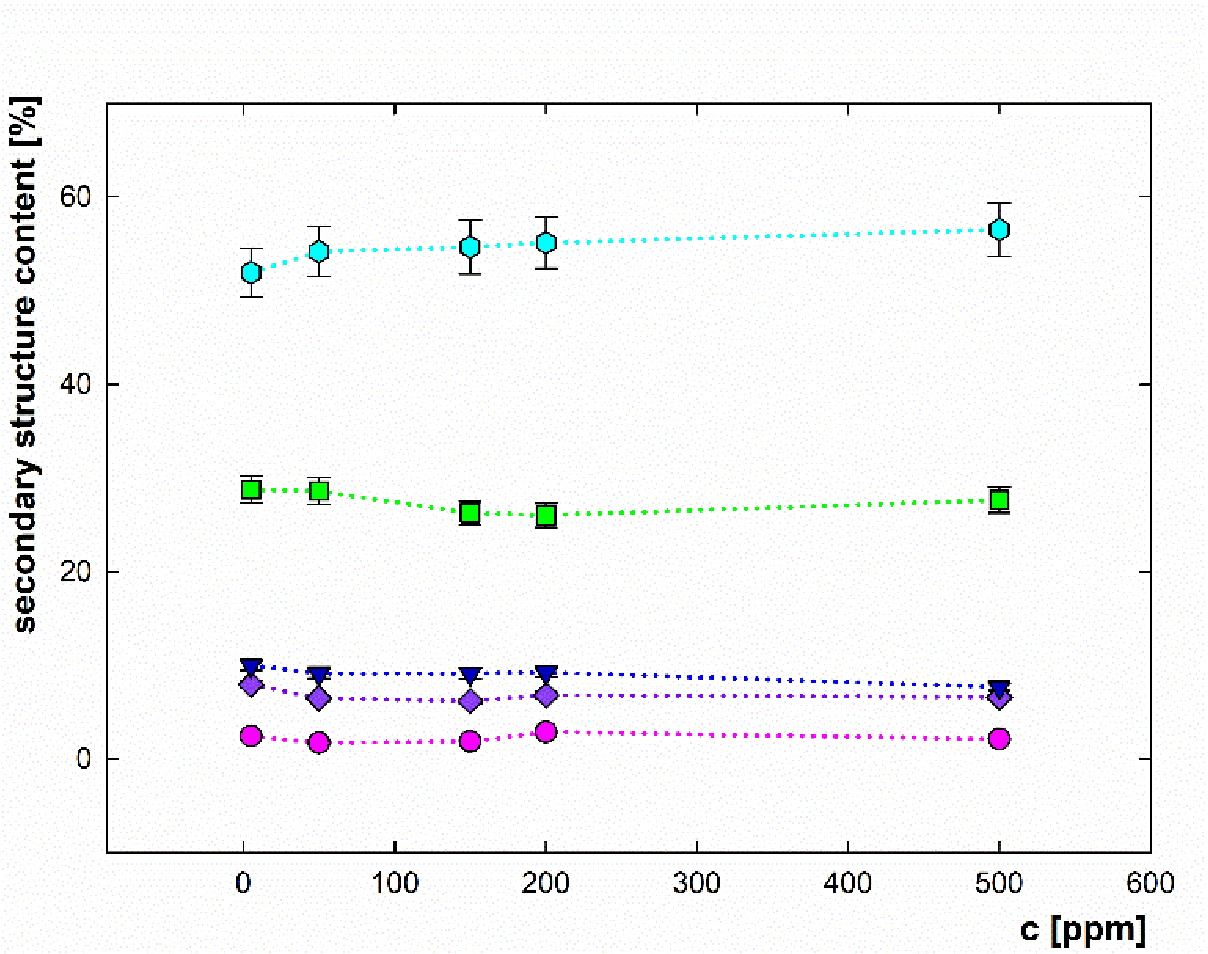


Figure SM7. The content of the BSA secondary structure components adsorbed on the gold surface depending on the solution concentration at I = 0.01 M NaCl, pH = 6.0: α-helices (cyan, ⬢), β-turns (green, ▪), disordered structures ( navy blue, ▲), β-sheets (purple, ◆), $3_{10}$-helix (pink, ●).
